# Insights into the structure-function relationship of the NorQ/NorD chaperones from *Paracoccus denitrificans* reveal shared principles of interacting MoxR AAA+/VWA domain proteins

**DOI:** 10.1101/2022.11.16.516607

**Authors:** Maximilian Kahle, Sofia Appelgren, Arne Elofsson, Marta Carroni, Pia Ädelroth

**Affiliations:** Department of Biochemistry and Biophysics, Stockholm University, SE-106 91 Stockholm, Sweden; Swedish Cryo-EM Facility, Science for Life Laboratory Stockholm University, Solna, Sweden; Department of Biochemistry, University of Potsdam, 14476 Potsdam, Germany

**Author notes:** Authors contributed equally (Maximilian Kahle, Sofia Appelgren). Address correspondence to: Pia Ädelroth. Maximilian Kahle. Marta Carroni.

**Keywords:** iron, nitric oxide reductase, *c*NOR, VWA, AAA+, ATPase, Fe_B_, protein remodeling, *nor* accessory genes, MoxR, cryo-EM, AlphaFold

## Abstract

NorQ, a member of the MoxR-class of AAA+ ATPases, and NorD, a protein containing a Von Willebrand Factor Type A (VWA) domain, are essential for non-heme iron (Fe_B_) cofactor insertion into cytochrome *c*-dependent nitric oxide reductase (*c*NOR). *c*NOR catalyzes the NO reduction, a key step of bacterial denitrification. This work aimed at elucidating the specific mechanism of NorQD-catalyzed Fe_B_ insertion, and the general mechanism of the MoxR/VWA interacting protein families. We show that NorQ-catalyzed ATP hydrolysis, an intact VWA-domain in NorD and specific surface carboxylates on *c*NOR are all features required for *c*NOR activation. Supported by BN-PAGE, low-resolution cryo-EM structures of NorQ and the NorQD complex show that NorQ forms a circular hexamer with a monomer of NorD binding both to the side and to the central pore of the NorQ ring. Guided by AlphaFold predictions, we assign the density that ‘plugs’ the NorQ ring pore to the VWA domain of NorD with a protruding ‘finger’ inserting through the pore, and suggest this binding mode to be general for MoxR/VWA couples. We present a tentative model for the mechanism of NorQD-catalyzed *c*NOR remodelling and suggest many of its features to be applicable to the whole MoxR/VWA family.

## INTRODUCTION

Metalloproteins constitute at least 30% of all known proteins and are used for a multitude of important functions in the cell, such as oxygen binding and transport, catalysis and electron-transfer chemistry. In many cases, the metal cofactors are inserted by dedicated chaperones during assembly of the metalloproteins (1). The bacterial MoxR proteins are involved in a variety of such cofactor assembly/re-organization processes and their detailed mechanism of action is poorly understood. We recently showed that the MoxR protein NorQ from *Paracoccus (P*.*) denitrificans* is, together with NorD, its partner protein, essential for the insertion of the Fe_B_ cofactor into the nitric oxide reductase (NOR) protein (2). Cytochrome (cyt.) *c*-dependent nitric oxide reductase (*c*NOR) is a two-subunit (NorCB) membrane protein complex, which catalyzes the reduction of toxic NO to N_2_O (2NO + 2e^−^ + 2H^+^ → N_2_O + H_2_O). This enzyme forms part of the bacterial denitrification chain, which involves the stepwise reduction of NO_3-_ to N_2_. *c*NOR is related to the terminal oxidases of aerobic respiratory chains, including mitochondrial cytochrome *c* oxidase, as it belongs to the same superfamily of heme-copper oxidases (HCuOs) (3, 4), where *c*NORs form their own subfamily. The catalytic subunit NorB is an integral membrane protein that contains a low-spin heme *b*, a high-spin heme *b*_3_ and a non-heme iron Fe_B_. Heme *b*_3_ and Fe_B_ form the binuclear active site and are spin-coupled via a μ-oxo bridge (5–7). The NorC subunit, which is a *c*-cytochrome, is membrane-anchored by one helix and has a soluble domain in the periplasm. The *c*NOR structure also contains calcium, bound at the interface of NorB and NorC (5). Protons and electrons needed for NO reduction are taken up from the periplasm, which is why the reaction catalyzed by *c*NOR is non-electrogenic (8, 9), in contrast to the proton-pumping O_2_-reducing HCuOs.

A set of specific chaperones is needed for assembly of the O_2_-reducing HCuOs. This includes the insertion of heme groups and copper cofactors, e.g. copper insertion into the *Rhodobacter (R*.*) sphaeroides aa*_3_ oxidase (an A-type HCuO, for a description of subfamilies, see (10)) is facilitated by PCuAC, PrrC and Cox11, and heme *a*_3_ insertion depends on Surf1p (11, 12). In *R. capsulatus*, the insertion of Cu_B_ into *cbb*_3_ oxidase (a C-type HCuO) was shown to rely on similar copper-binding proteins (13). However, only little is known about the assembly and cofactor insertion process of *c*NOR.

The *nor* operon of *Paracoccus* (*P*.) *denitrificans* comprises six genes (*norCBQDEF)* (14), but only *norC* and *norB* are coding for proteins present in the *c*NOR complex (5, 15, 16). Mutational analysis supports the idea that *norQ* and *norD* are cooperatively involved in activation of *c*NOR on the protein level (14, 17, 18). Recently, we reported that *c*NOR expressed in the absence of *norQDEF* forms a stable NorCB dimer that is fully hemylated but lacks the Fe_B_ cofactor and is therefore inactive. We further showed that the NorQ and NorD specifically act as chaperones involved in Fe_B_ insertion (2, 19). In contrast, the homologous quinol-oxidizing qNOR, expressed from a single gene *norZ*, does not require any additional genes for Fe_B_ insertion and activity (17).

NorQ is a member of the large MoxR protein subfamily of bacterial AAA+ ATPases (20, 21). AAA+ ATPases have diverse functions in the cell and are present in all three domains of life (22, 23). AAA+ proteins are molecular motors that typically assemble as hexameric rings and share a function in remodeling and/or unfolding macromolecules. AAA+ proteins were classified into seven different clades based on evolutionary lineage and structural insertions (see (23, 24) for recent reviews), where the MoxR proteins belong to clade 7 which also contains e.g. the dynein families and has the helix 2 insert (H2I). Recently though, this classification was challenged (24) since the cryo-EM ‘boom’ in structural information shows that e.g. the hexameric CbbQ, a clade 7 MoxR protein closely related to NorQ, structurally resembles the clade 6 McrB hexamer involved in DNA cleavage (25, 26) more than it resembles the hexameric assembly of the MoxR protein RavA (27). The MoxR AAA+ proteins are among the least studied families of AAA+ proteins, but it has been speculated that they primarily act as molecular chaperones involved in metal cofactor insertion or other remodeling events (21).

NorQ (Fig. 1) consists of a single AAA+ module, which contains several conserved motifs, such as the Walker A and Walker B (WB) motif, which promote nucleotide binding and hydrolysis, respectively. The Walker A motif contains a critical lysine that stabilizes ATP binding, and the Walker B motif contains a conserved aspartate for binding of Mg^2+^ followed by a catalytic glutamate essential for ATP hydrolysis. In most AAA+ proteins, the ATP binding site is located at the interface between protomers with a conserved Arg finger acting in *trans*, making the AAA+ ATPase functional only in its multimeric state. The mechanistic sequence of AAA+ protein function is not fully understood but often involves an asymmetric organization of protomers in different nucleotide-bound states in the ring oligomer (28).

**Figure 1.**
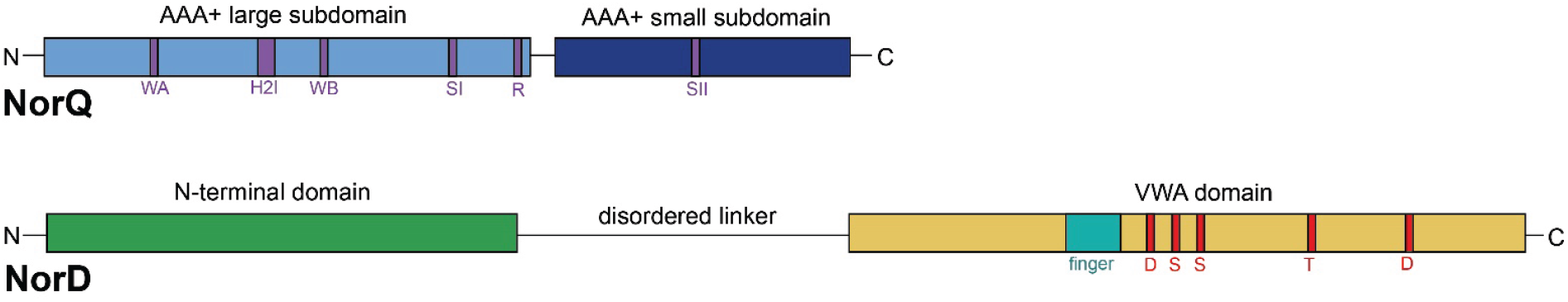
Schematic domain architecture of NorQ and NorD. NorQ is a single AAA+ ATPase module that consists of the large α/β subdomain (light blue) and the C-terminal all-α subdomain (dark blue). The conserved motifs of the AAA+ module are marked (purple): Walker A (WA), H2I (helix 2 insert), Walker B (WB), sensor I (SI), arginine finger (R) and sensor II (SII). NorD consists of a N-terminal domain (green) and a C-terminal Von Willebrand Factor A domain (yellow), connected via a disordered linker (see text). The conserved metal ion-dependent adhesion site (MIDAS) residues (red) are marked. For the predicted “finger”, see text.

NorD (Fig. 1) is a 70 kDa protein that contains a 33 kDa C-terminal Von Willebrand Factor Type A (VWA) domain. VWA domains are believed to generally function by mediating metal ion-dependent protein-protein interactions (29). Within the VWA domain the MIDAS (metal ion-dependent adhesion site) motif (DxSxS-T-D) is involved in coordinating a divalent metal ion (often Mg^2+^). A sixth ligand of the coordination sphere is an acidic surface residue donated by the interacting target protein, thereby stabilizing the protein-protein interaction (29). The protein sequence preceding the VWA domain in NorD shows no obvious homology to any other proteins in the databases. NorQ and NorD are most likely cytosolic proteins since they lack signal peptides.

Often, MoxR proteins genetically pair with VWA-carrying proteins (as for *norQD*), suggesting a functional relation (20). MoxR AAA+/VWA interactions have been established for the RavA/ViaA couple interacting with fumarate reductase in *E. coli* (30) and the CbbQ/CbbO couple involved in Rubisco activation in multiple bacteria (31), where ViaA and CbbO carry C-terminal VWA domains.

In this work we established the expression and purification of the NorQ and the NorQD complex and investigated their functional and structural properties. We also identified the binding site for the NorQD complex on the surface of NorB target protein. Our data shed light on the Fe_B_-insertion mechanism for *c*NOR activation specifically and the structure-function relationships in the MoxR proteins and their binding to VWA-containing partner proteins in general.

## RESULTS

### Expression and purification of the NorQ and NorD proteins

The wildtype 6-His-tagged NorQ protein was purified from *E. coli* (Fig. 2a) at a yield of ~40 mg/L culture. The Walker A (WA) variant of NorQ (K48A) was expressed but prone to precipitation and was not purified. The Walker B (WB) variant of NorQ (NorQ^WB^, E109Q) expressed similarly to wildtype.

**Figure 2:**
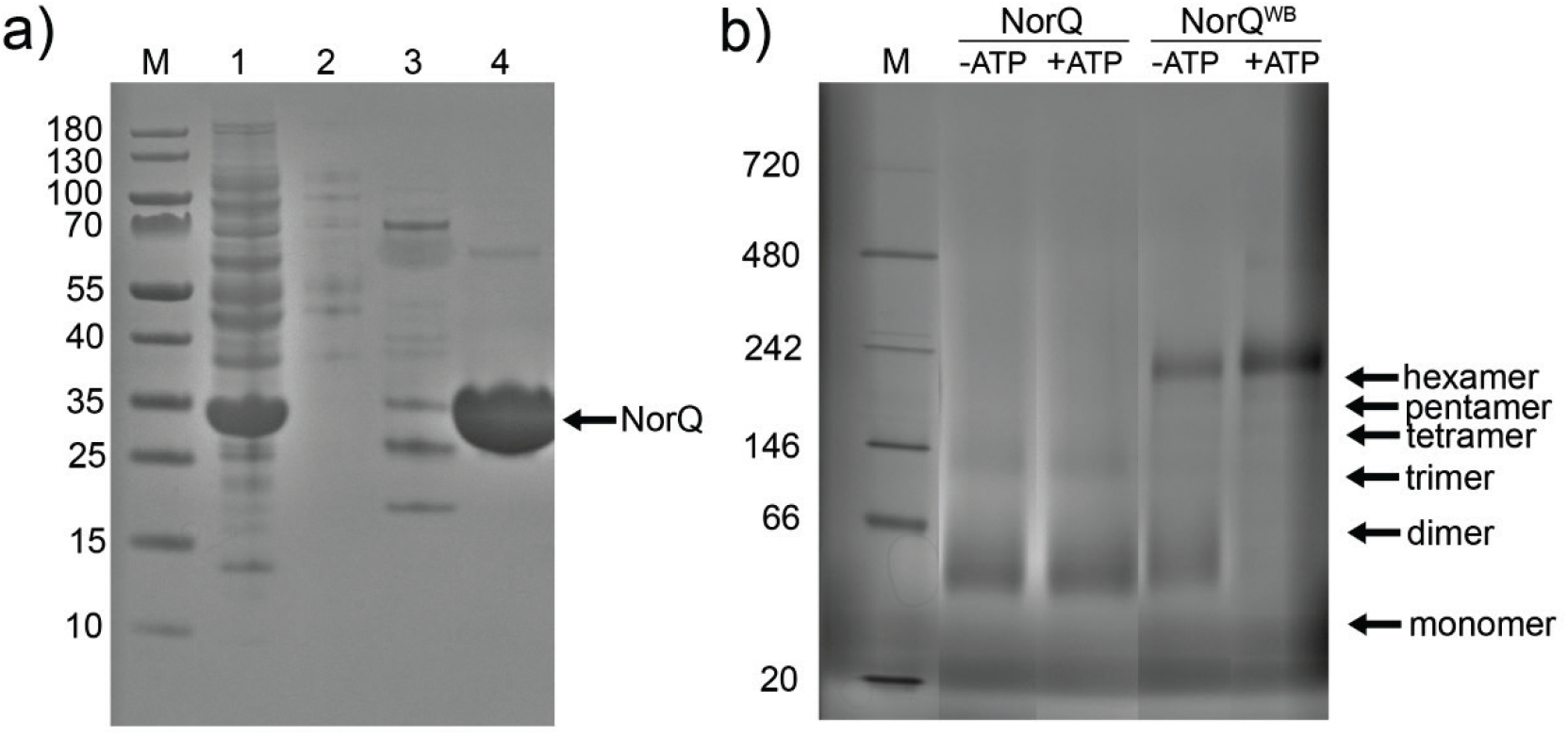
SDS-PAGE and Blue Native-PAGE for NorQ and NorQ^WB^. a) SDS-PAGE of the purification process of NorQ. Experimental details: Gel run at 100 V. Lane 1: Cytosolic suspension. Lane 2: flow-through from Ni-NTA. Lane 3: Wash with 150-250 mM imidazole. Lane 4: Elution with a 250-500 mM imidazole gradient. b) BN-PAGE of NorQ (14 μg) and NorQ^WB^ (18 μg), supplemented with 2 mM ATP and 20 mM MgCl_2_. Gel run for 1h at 150 V and 45 min at 250 V. The expected locations of several oligomeric states of NorQ are indicated. Original gel in Fig. S1a.

BN-PAGE and SEC analysis (Fig. 2b, Fig. S1, Fig. S2) showed that the NorQ protein forms multiple oligomeric states with sizes ranging from 30 kDa (monomer) to ~180 kDa (hexamer, more clearly observed in the NorQ^WB^, Fig. 2b and Fig S1a) with trimers, tetramers and pentamers also clearly present. The addition of ATP to the BN-PAGE experiments showed a clear shifting of the oligomerization state to the hexamer (Fig. 2b, and Fig. S1a) for the NorQ^WB^, linked to its loss of ATPase activity (see below).

The NorD protein could not be expressed in the absence of NorQ, most likely due to low solubility of the protein. Co-expression of 6-His-tagged NorD and untagged NorQ led to purification of the NorQD via affinity chromatography (Fig. 3a). The identity of NorQ and NorD on SDS-PAGE was confirmed by mass spectrometry analysis of excised protein bands (see Fig. S3, Table S1).

**Figure 3:**
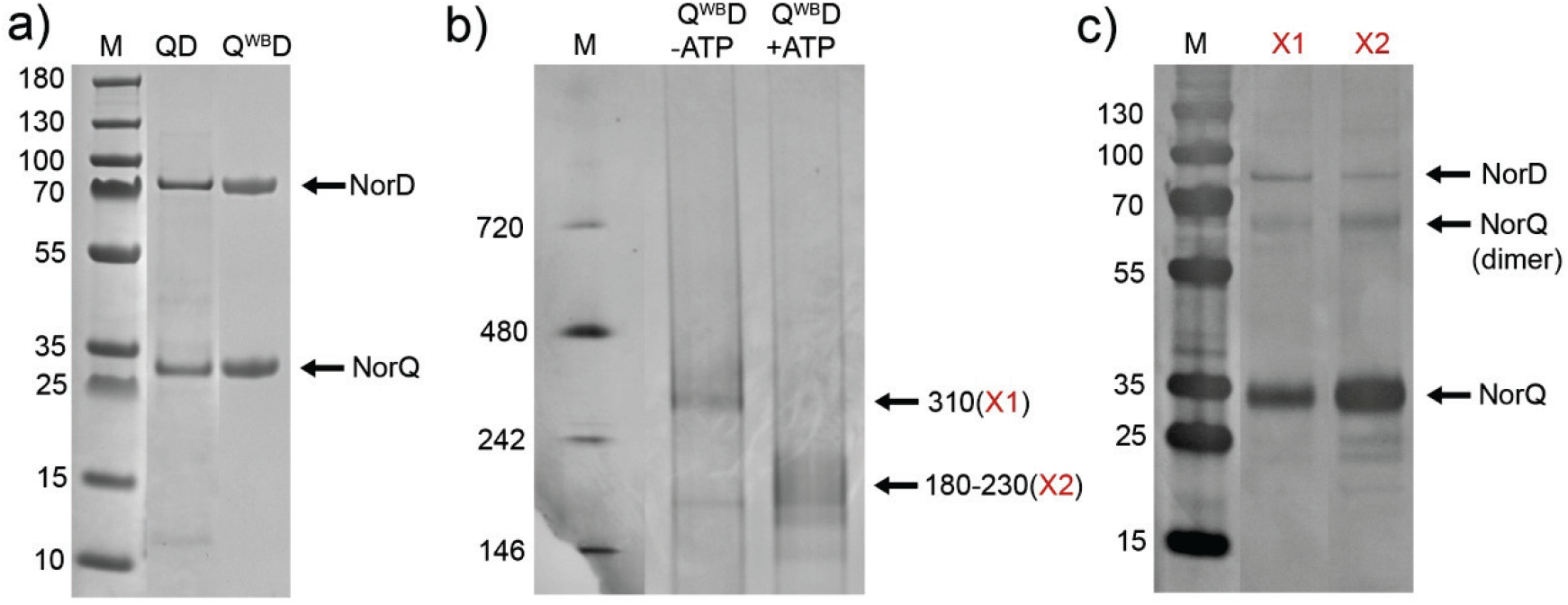
SDS-PAGE and Blue Native-PAGE for NorQD and NorQ^WB^D. a) SDS PAGE of gradient-purified NorQD and NorQ^WB^D. ~2 μg protein was loaded on the gel and run at 100 V. b) Blue Native PAGE of NorQ^WB^D. ~10 μg protein was loaded and the gel was run for 1h at 150 V and 45 min at 250 V. c) Second dimension SDS-PAGE of individual bands of the Blue-Native-PAGE gel (panel b). The bands were visualized by silver staining and are labeled as identified by mass spectrometry (see Table S1 and Fig. S3e). Original gels in Fig. S1 and Fig. S3.

The purified protein complex was prone to precipitate when concentrated, when kept at temperatures > 4 °C or with increasing number of freeze/thaw cycles. This is possibly related to the intrinsic property of NorQ to oligomerize and of the VWA domain in NorD to promote metal ion-dependent interactions. Proteins homologous to NorQ and NorD were also found prone to aggregate (31, 32).The NorQ^WB^D complex was generally better expressed and less prone to precipitation than the WT NorQD complex and band patterns on SDS-PAGE were very similar for both (Fig. 3a). BN-PAGE and SEC analysis (Fig. 3b, Fig. S1b, and Fig. S2b) showed that the NorQD proteins form multiple oligomeric states with sizes ranging from ~300 kDa, to ~30 kDa (presumably monomeric NorQ). Band size distribution shifted in the presence of ATP (Fig. 3b, Fig. S1b and Fig. S2b).

A second dimension SDS-PAGE for bands excised from the BN-PAGE and mass-spec analysis (Fig. 3c, Fig. S3, Table S1) showed that the 310 kDa band in the NorQ^WB^D preparation most clearly contained both NorD (70 kDa) and NorQ (30 kDa). This band size is consistent with a hexameric NorQ (6×30 kDa) bound to a monomer (or possibly dimer) of NorD (1×70 kDa). After addition of ATP this band distributes to lower molecular weights (180-230 kDa) consistent with a trimer to hexamer of NorQ bound to a monomer of NorD (Fig. 3b).

Mutation of the conserved MIDAS residues in the VWA domain of NorD (T534V or D562N, see sequence alignment in Fig. S4) still allowed co-purification of NorQ (Fig. S3d), indicating that the NorQ-NorD interaction site does not involve this protein region.

### Single particle cryo-EM

Because of the higher solubility of NorQ^WB^, this variant was used for all cryo-EM work. 2D classification of NorQ^WB^ included about 42,000 particles (from a total number of about 130,000 picked particles) and showed various views (Fig. S5). Initial models were calculated with and without 6-fold symmetry applied (Figure 4a). The C1 *ab initio* models showed open ring and asymmetric ring assemblies of NorQ^WB^. After 3D classification, refinement yielded two maps, one processed always in C1 and one processed in C6, but for which symmetry was relaxed to C1 at later stages (Figure 4b). 3D maps processed in C6 tended to give a higher nominal resolution (~8 Å) compared with models without symmetry constraints (~11 Å) but not enough to justify the existence of a fully 6-fold symmetric assembly.

**Figure 4:**
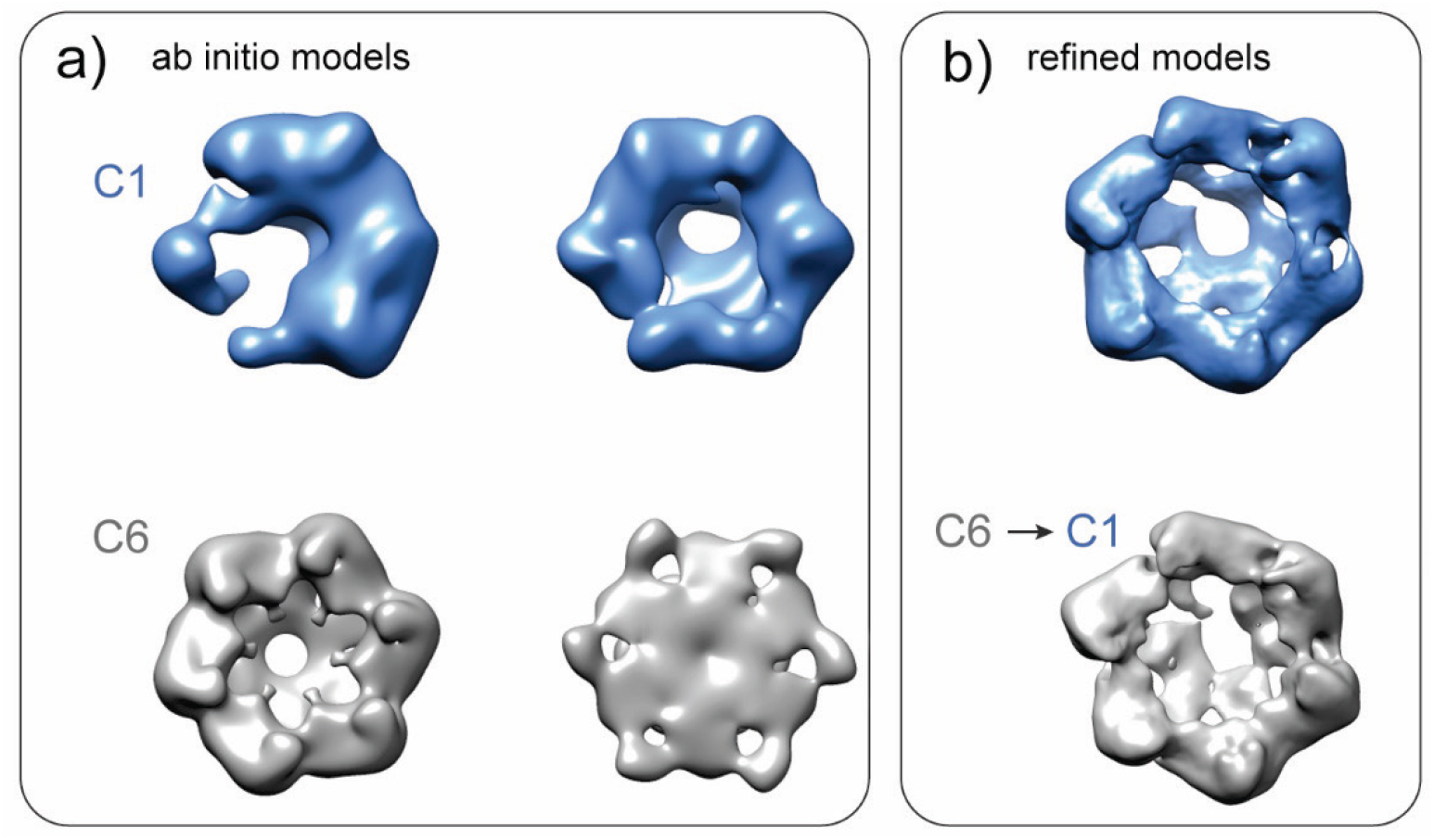
Ab initio models (a) and refined models (b) of NorQ^WB^. a) Two different ab initio models in C1 (blue models) and another two initial models in C6 (grey models) are shown. b) A refined model processed in C1 is shown in blue. Another model which was refined using C6 constraints and then symmetry relaxed to C1 is shown in grey.

Refinement of a C6 structure subpopulation of NorQ^WB^ (~30,000 particles) yielded a nominal resolution of ~8 Å (Figure 5). A rigid body fit of the crystal structure of the homologous csoCbbQ from *Halothiobacillus (H*.*) neapolitanus* (PDB: 5C3C (33), 49% sequence identity) as well as the recently solved structure of AfCbbQ2 from *Acidithiobacillus ferrooxidans* (25) with 53% sequence identity (PDB: 6L1Q ((25)) was found to match the 3D map of NorQ^WB^ (Figure 5) relatively well. The recent structure of CbbQ (PDB ID: 6L1Q) resolved for the first time the H2I region (residue 79-93 in the alignment in Fig. S6a) of CbbQ, and this region is resolved also partially in our 8Å NorQ structure (Fig. 5b). The residues discussed in the text as well as the monomer/monomer interaction surface in the NorQ hexamer are highlighted in the NorQ homology model in Fig. S6bc.

**Figure 5:**
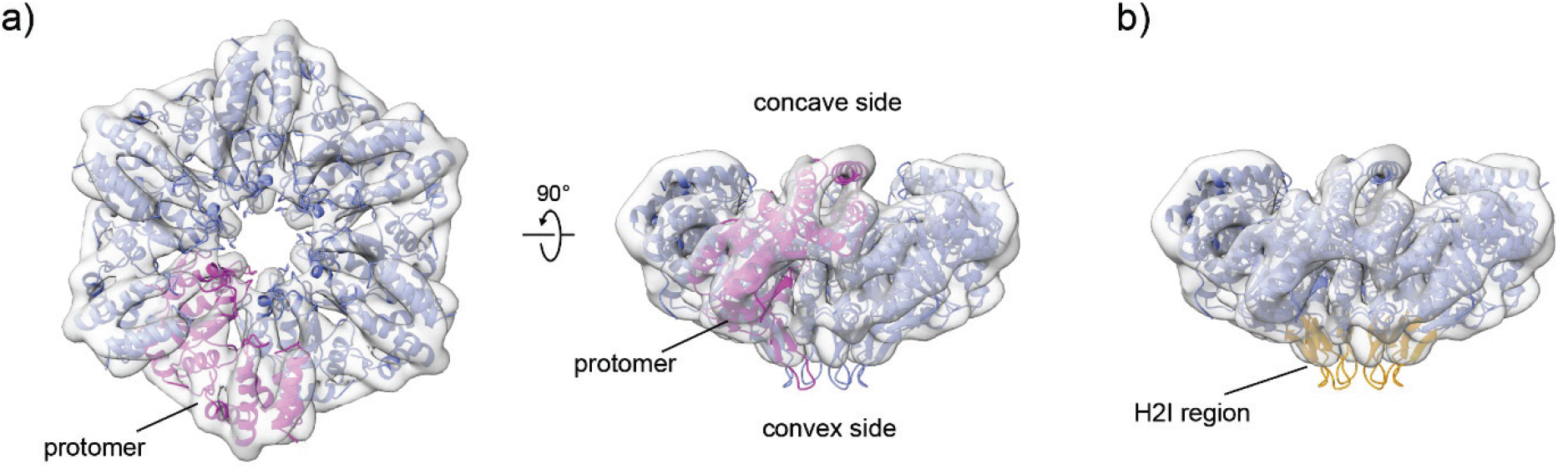
Refined 3D model of NorQ^WB^. The cryo-EM map at ~8 Å (grey surface) and the rigid body fit of the NorQ homology model based on the AfCbbQ2 crystal structure (PDB: 6L1Q (25)) is shown. a) Within the NorQ hexamer (blue) one protomer is highlighted in pink. b) The H2Is from all subunits protruding at the convex side of the hexamer are shown in yellow.

We further investigated the NorQ^WB^ in complex with NorD to gather additional information about the stoichiometry of the functional NorQ^WB^D. 2D class averages from a subset of about 500,000 clean particles of NorQ^WB^D were found to be consistent with a hexameric, ring-shaped oligomer. Most classes show an additional ‘comma-like’ density in close proximity to the ring (Fig. 6a). Because of strong preferred particle orientation, while up to 3Å resolution was achievable in one direction it was possible to determine an isotropic map only at 10Å. This low-resolution 3D map shows the presence of an asymmetric hexamer of NorQ^WB^ with density for NorD acting like a plug inserted in the middle of the hexameric ring (Fig. 6b). The ‘comma’ to the side of the ring was not present in the refined model, presumably because of flexibility or conformational variability (see below and Discussion).

**Figure 6:**
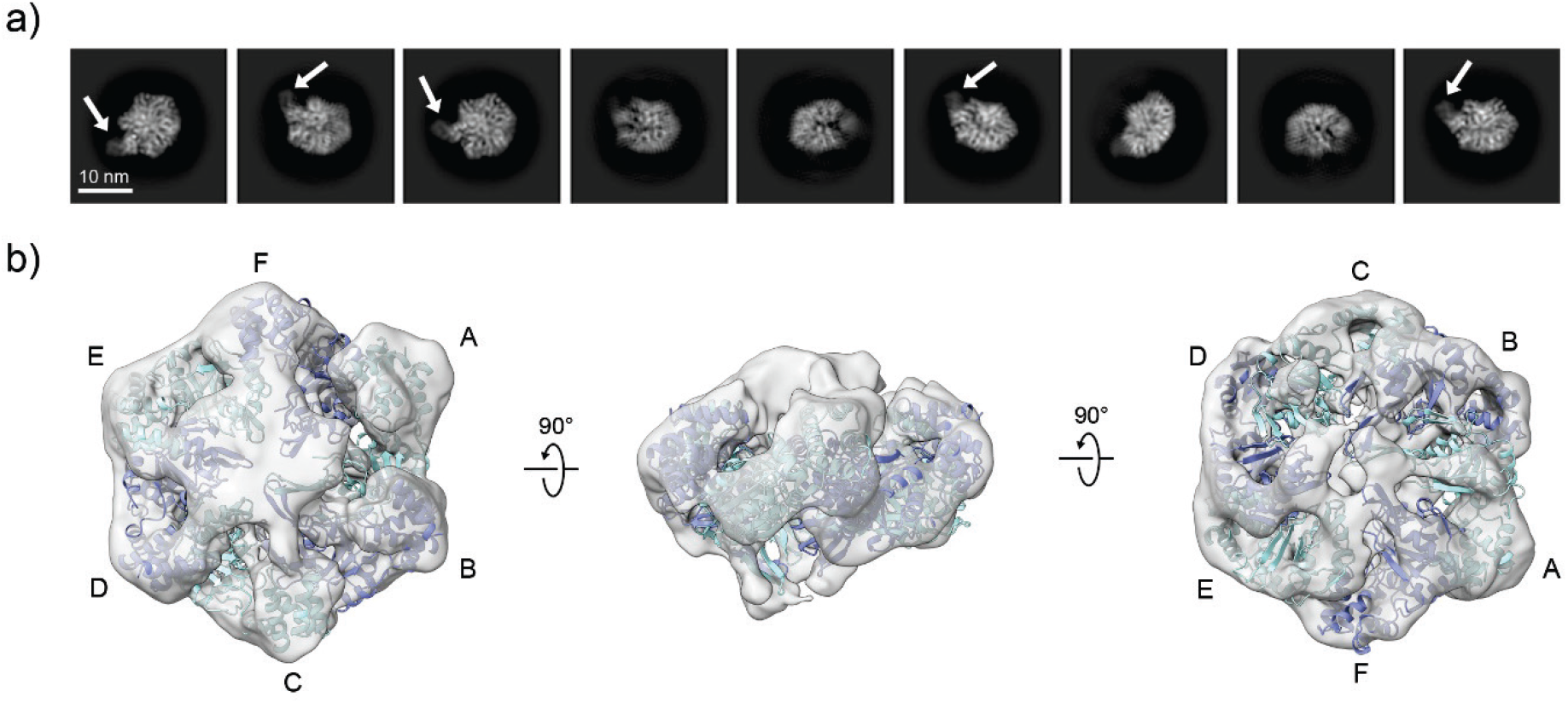
NorQ^WB^D complex in cryo-EM. Cryo-EM samples contained 1 mg/mL protein in 10 mM TRIS/HCl, pH 8, 150 mM NaCl, 10% glycerol. a) Representative 2D class averages, with the ‘comma’ visible to the side of the ring marked with arrows. b) Refined cryo-EM 3D-map at ~10 Å. Shown is also the modelled NorQ hexameric ring with the protomers labelled A-F (see text).

### Model building

In order to fit the extra density to NorD, we first predicted its structure using the recently developed tool AlphaFold (34) (AF), see Fig. 7a. The predicted structure of NorD shows two well folded domains separated by a non-structured region, also predicted by conventional tools (see Fig. S7) to be disordered. The prediction also shows a new feature not observed nor modeled previously in the VWA domain of NorD or in other VWA-domain containing proteins that partner with MoxR AAA+ proteins (Fig. S8). This new feature is a ‘finger’, stabilized by a β-sheet, that sticks out of the VWA globular domain. We suggest this ‘finger’ binds in the hexameric AAA+ central pore and positions the MIDAS residues of the VWA domain ‘on top’ of the ring (see below and Discussion).

**Figure 7:**
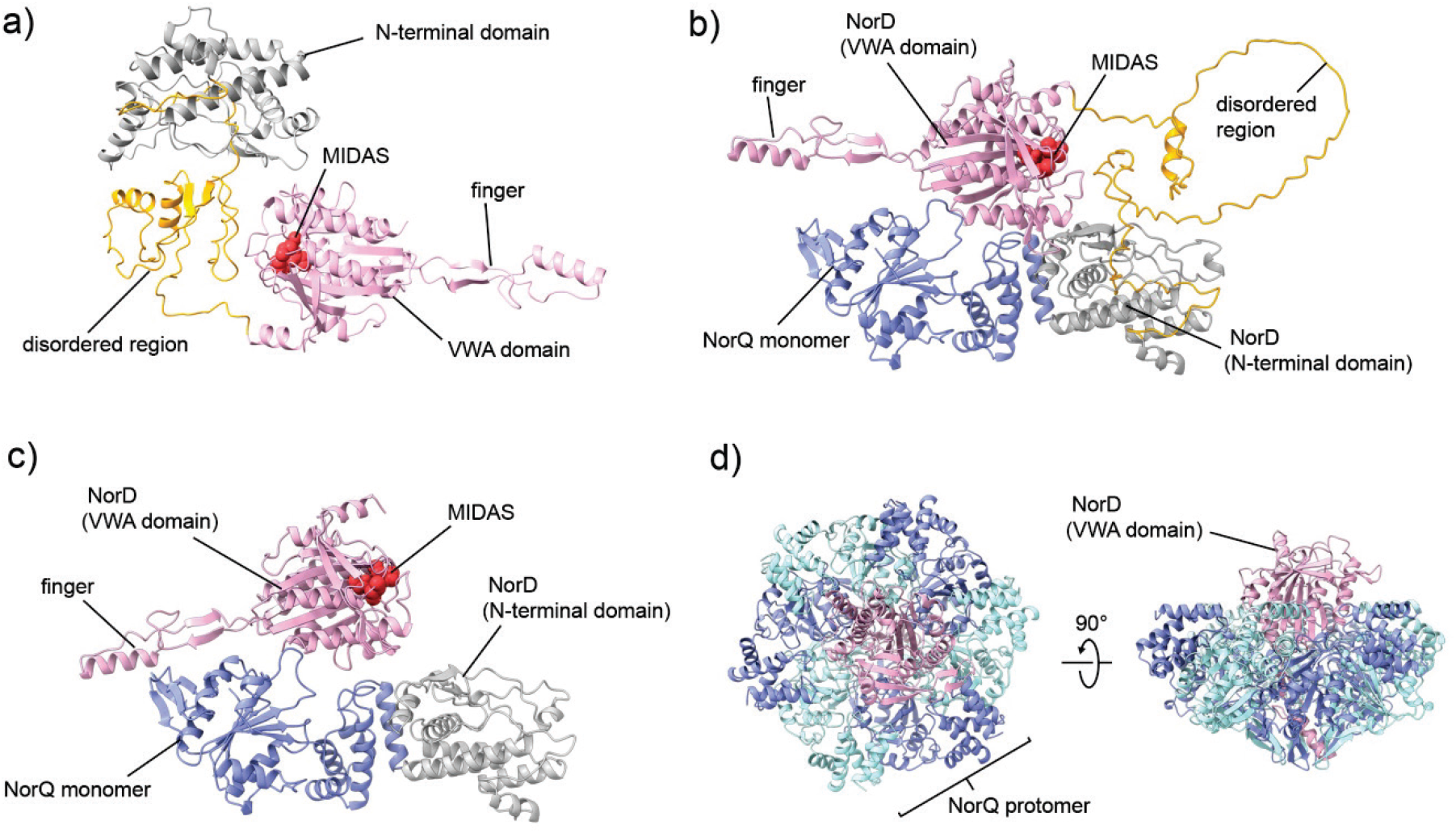
Single chain and multimer models of NorQ and NorD using AlphaFold. a) Prediction of full-length NorD (Uniprot: Q51665) showing the N-terminal domain (grey), disordered region (yellow) and VWA domain (pink) with the MIDAS residues in red. b) Multimer model of one NorD and one NorQ (blue, Uniprot: Q51663). Coloration of NorD is as in a). c) Multimer model of one NorQ, the N-terminal domain of NorD (grey, residues 1-224) and the NorD VWA-domain (pink, 336-638, MIDAS in red). d) Multimer prediction with six chains of NorQ (alternating blue and cyan) and one NorD VWA domain (pink).

For different version of a complex between NorQ and NorD, we used AlphaFold-multimer (35). We built a complete complex of 6 NorQ chains and one NorD chain. This complex consistently put the VWA domain of NorD in the center of a hexameric ring of NorQ (see Fig. S9).The predicted TM score (pTM, see Methods) for this model is 0.41, indicating that it is of limited confidence. (see Table S2 for pTM scores for all models shown). We also modeled a complex of the 6 NorQ monomers (forming a hexamer) with only the VWA domain of NorD (Fig. 7d). The pTM for these models are 0.45-0.52, somewhat higher than for the full complex. To obtain a better picture of the interaction between NorD and NorQ we also modeled a NorQ monomer-NorD monomer complex (Fig. 7b, pTM significantly higher at 0.7). We also modeled this complex without the disordered domain of NorD, i.e. the dimer was modeled as a trimer with the NorD N-terminal domain, the NorD (C-terminal) VWA domain and the entire NorQ chain (Fig. 7c, pTM-scores 0.63-0.74, similar to the full monomer-monomer complex). This model was then used for fitting into the EM density.

We generated a hexameric NorQ model to fit into the cryo-EM density (Fig. 6b) based on the hexameric asymmetric structure of the AAA+ protein ClpX (PDB:6SFW) (36), by superimposing the AF NorQ monomer onto each of the ClpX protomers. The rationale behind is the rough placement of the NorQ protomers in the AAA+ ring with the seam subunit (protomer A) displaced (Fig. 6b). Even at this resolution it is in fact clear that the NorQ ring is not perfectly symmetric similarly to many other AAA+ structures (see e.g. also the McrBC complex (PDB: 6UT5 and 6UT6) (26, 37)). We assume that the observed density plugged into the NorQ ring is the VWA domain of NorD, based also on the observation of analogous low-resolution densities for the MoxR-VWA complex CbbQ-CbbO (25, 31) where the VWA domain was identified as the density bound to the pore of the ring. The NorQ-NorD AF model (Fig. 7b, omitting the N-terminal domain of NorD) was thus overlaid onto the NorQ ring protomer that would give the least of clashes, thus obtaining a full 6NorQ-NorD-VWA domain model. This overall binding topology with the VWA domain of NorD inserted into the NorQ pore, was also supported by the AF prediction shown in Fig. 7d. The 6NorQ-NorD model was flexibly fitted into the cryo-EM map using iMODFIT (38). The resulting 6NorQ-NorD model (Fig. 8abc and see Fig. S10 for comparison to the NorQ_6_VWA model), qualitatively consistent with the data, does however not perfectly fit the density, especially at the convex side of the ring and there is ‘extra’ density that could be the VWA domain ‘finger’ or the NorQ H2I sitting in different positions (see Fig. S11 and Discussion).

**Figure 8:**
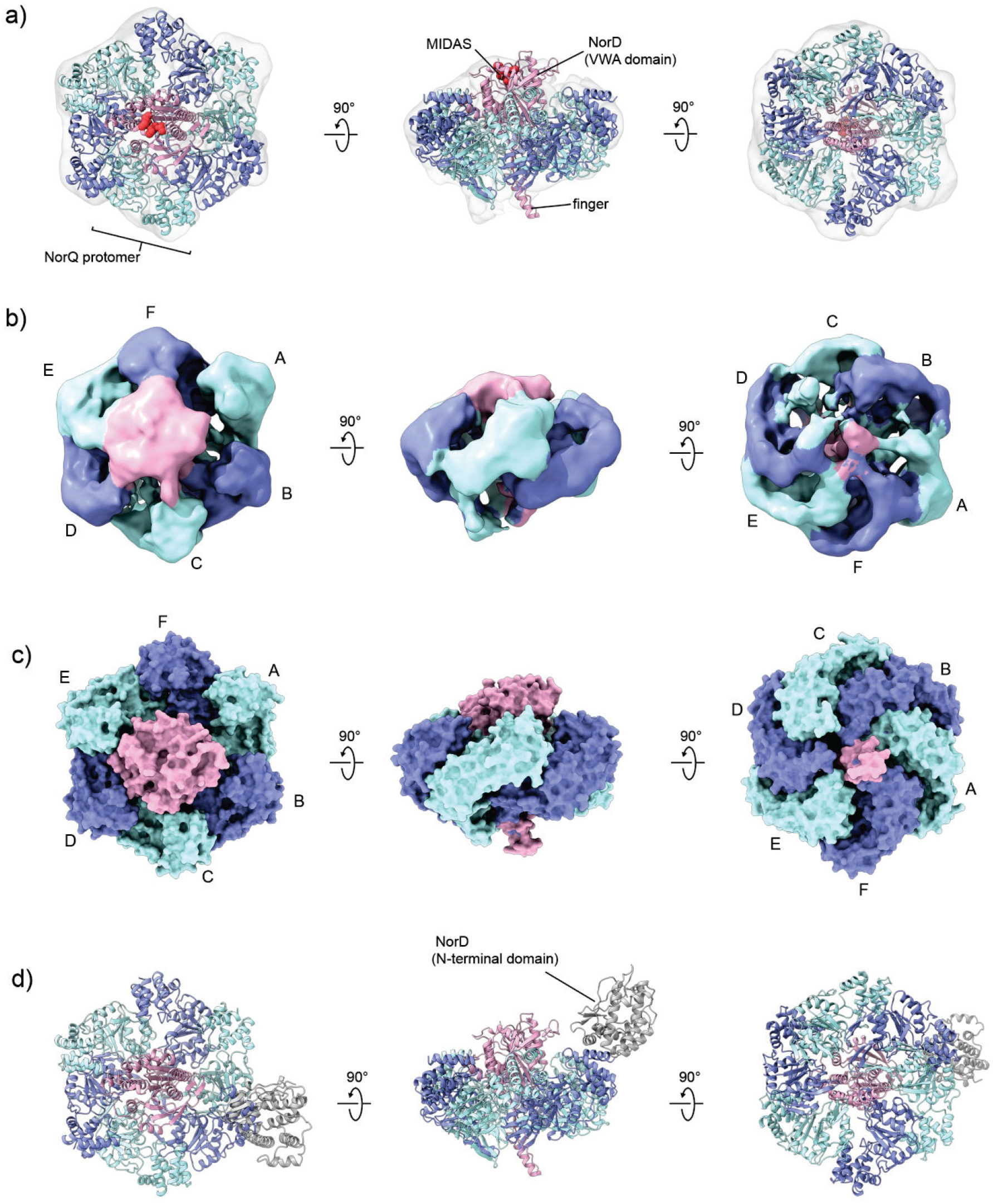
Models of the NorQ^WB^D complex with combined data from cryo-EM analysis and AlphaFold predictions. The NorQ hexamer is shown in blue and cyan, the NorD VWA domain in pink with MIDAS residues in red, and the N-terminal domain in grey (d). a) Model of the NorQ hexamer bound to the NorD VWA domain based on the AF prediction shown in Fig. 7c and after flexible fitting into the cryo-EM map (Fig. 6b). b) Cryo-EM map colored according to the fitted NorQD model shown in a) and with the NorQ protomers labelled as in Fig. 6b) Surface representation of the fitted NorQD model. d) The NorD N-terminal domain was added to the fitted NorQD model preserving its relative position as predicted (Fig. 7c).

We further assume that the N-terminal of NorD is located to the side of the NorQ hexamer and is represented by the ‘comma’ visible in the 2D classes (Fig. 6a) but is missing in the refined density. Fig. 8d shows the tentative location for the NorD N-terminal using the NorQ-NorD AF prediction (Fig. 7c). The overall orientation of the NorD domains relative to each other and to the NorQ hexamer as modeled in Fig. 7b and Fig.7c was supported by the AF prediction of the complex with the entire NorD protein (Fig. S9).

### ATPase activity of NorQ and the NorQD complex

The purified NorQ exhibited ‘uncoupled’ ATP hydrolysis activity of ~100 nmol mg^−1^ min^−1^, measured with both the pyruvate kinase coupled assay (Table 1) and the malachite green ‘end-point’ assay (Table 1 and Fig. S12). This corresponds to a *k*_cat_ of 25/(min*hexameric complex) and the *K*_m_ was measured to 0.16±0.02 mM ATP (Fig. S12). This activity was lost in the NorQ^WB^ variant (Table 1, Fig. S12).

**Table 1:** ATPase activity of purified NorQ and NorQD complex constructs. Experimental conditions for the NADH-coupled assay: 50 mM TRIS/HCl, pH 8.0, 150 mM NaCl, 15 mM MgCl_2_, 2.5 mM ATP, 1 mM phosphoenolpyruvate, 0.3 mM NADH, 12 U/mL pyruvate kinase, 12 U/mL lactate dehydrogenase. Experimental conditions for the malachite green assay: 50 mM TRIS/HCl, pH 8, 150 mM NaCl, 10% (v/v) glycerol, 20 mM MgCl_2_, 2 mM ATP. Data presented as averages with standard deviation. dt=double-tagged construct.

| Protein construct | malachite green assay<br>(nmol mg<sup>-1</sup> min<sup>-1</sup>) | NADH-coupled assay<br>(nmol mg<sup>-1</sup> min<sup>-1</sup>) | NADH-coupled assay<br>(min<sup>-1</sup>/complex) |
| --- | --- | --- | --- |
| NorQ | 72 ± 2 (100%) (n=2) | 141 ± (100%) (n=3) | 25 ± 2 (100%) (n=3) |
| NorQ<sup>WB</sup> | 2.0 ± 0.3 (3%) (n=2) | 1.5 ± 0.3 (1%) (n=3) | 0.30 ± 0.05 (1%) (n=3) |
| NorQD | 9 ± 2 (13%) (n=2) | 30 ± 1.3 (21%) (n=3) | 7.4 ± 0.3 (30%) (n=3) |
| NorQ<sup>WB</sup>D | 7 ± 1 (10%) (n=2) | 13 ± 0.4 (9%) (n=3) | 3.2 ± 0.1 (13%) (n=3) |
| dt-NorQD | 3.8 ± 1.3 (5%) (n=2) | 2.9 ± 2.7 (2%) (n=2) | 0.7 ± 0.6 (3%) (n=2) |
| dt-NorQ<sup>WB</sup>D | 1.3 ± 0.4 (2%) (n=2) | 0.9 ± 0.8 (1%) (n=3) | 0.2 ± 0.2 (1%) (n=3) |

In the wildtype NorQNorD (QD) complex, the ATPase activity was ~10% of that in NorQ alone, only slightly higher than in the Q^WB^D preparation (Table 1), i.e. the ‘uncoupled’ activity of NorQ alone is higher than in the presence of NorD (see Discussion). Because the residual activity of the NorQ^WB^D preparation was higher than in the NorQ^WB^ alone, we engineered a double tag (dt) construct (see Methods) in order to improve the purity of the preparation such that we could exclude contaminant activity. The data (Table 1) on the double-tag ‘ultrapure’ NorQD complexes clearly show that ATPase activity is lost in the dt-NorQ^WB^D construct, and that the activity is higher in the wildtype NorQD. The wildtype NorQD complexes were more prone to aggregate under the measurement conditions such that the numbers for its activity could be somewhat underestimated (see Methods), but it is clear that the complexation with NorD inhibits NorQ activity significantly. Additions of the purified *c*NOR, either wildtype or the Fe_B_-less variant, had no effect on the ATPase activity.

### Binding of NorQD to NorB and insertion of the Fe_B_

We investigated the putative involvement of specific cytoplasmic surface residues of NorB as an interaction surface for the VWA domain of NorD by mutagenesis. Here, aspartate and glutamate residues that are exposed to the bulk (as judged from the *c*NOR crystal structure, PDB: 3O0R (5)) and conserved in *c*NOR but not in qNOR were considered as likely candidates for such interactions (see alignment in Fig. S13). This is because qNOR does not require homologs of NorQD for Fe_B_ insertion (17). Among four candidate residues, only mutation of D220 or E222 abolished *c*NOR activity completely (Table 2). The non-heme iron content was substantially decreased in the D220A (to 20 ± 2 %) and E222A (to 23 ± 3 %) *c*NOR variants compared to wildtype. In contrast, a double mutation of E75 and E78 on the same protein surface produced active *c*NOR (~50 % of WT activity).

**Table 2:** NOR activity and non-heme Fe content. NO-reduction activity was measured in detergent solubilized membranes. Experimental conditions: 25 nM cNOR in 50 mM Hepes, pH 7.5, 50 mM KCl, 1 % DDM, 10 μM horse heart cyt. c, 500 μM TMPD and 3 mM ascorbate at 22 °C. The non-heme Fe content was measured with purified cNOR using the ferene method. ^#^The data for WT and ΔnorQDEF was published previously (2). ND= not determined.

| ***nor* operon modification**<br>**(pNOREX-His)** | **cNOR activity (%)** | **cNOR non-heme Fe content (%)** |
| --- | --- | --- |
| WT | 100<sup>#</sup> | 100<sup>#</sup> |
| ΔnorQDEF | 0.4<sup>#</sup> | 20±7<sup>#</sup> |
| E109Q (NorQ) | 3±3 (n=2) | ND |
| T534V (NorD) | 2±2 (n=2) | ND |
| D562N (NorD) | 4±1 (n=2) | ND |
| E75A/E78A (NorB) | 40±20 (n=6) | ND |
| D220A (NorB) | 5±3 (n=2) | 20±2 (n=2) |
| E222A (NorB) | 4±2 (n=2) | 23±3 (n=2) |
| N-terminal His-tag (NorD) | 70±10 (n=3) | ND |
| N-terminal His-tag (NorQ) | 80±30 (n=3) | ND |

We further verified the functional importance of the WB-motif (E109) of NorQ and the MIDAS residues (T534 and D562) in NorD in the *c*NOR operon by the observation that exchanging them abolished the activity of the target protein *c*NOR (Table 2). His tags inserted at the N-terminus of NorD or NorQ were found not to interfere with NorQD function, as they caused only minor changes in *c*NOR activity (Table 2).

### Re-activation experiments with crude cell extracts

After expression of the *nor* operon in *E. coli*, membrane fractions and soluble fractions were mixed in different combinations according to Table S3 in order to test a possible re-activation effect on Fe_B_-less *c*NOR (membrane bound) by NorQ and NorD proteins (soluble). NorE and NorF proteins have no effect on heterologous *c*NOR expression (2). A drastic decrease in *c*NOR activity (to ~8 %) and non-heme iron content (to ~10 %) when NorQ and NorD were absent (*norCB(ΔQDEF)* samples) from both fractions (Table S3) is consistent with our earlier results (2). When membranes that contained Fe_B_-deficient *c*NOR were mixed with the soluble fraction of cells expressing the complete *nor* operon, purified *c*NOR showed a 2.5-fold increase in activity and a 2-fold increase in non-heme iron content. Mixing detergent solubilized Fe_B_-deficient *c*NOR with the purified NorQD did not recover *c*NOR activity.

## DISCUSSION

### NorQ is an AAA+ ATPase that forms hexameric rings

Purified NorQ forms several oligomeric states, but in the E109Q (NorQ^WB^, Walker B motif) NorQ variant in the presence of ATP (Fig. 2 and Fig. S1), the hexameric state dominates. This behavior is similar to that found for other MoxR and AAA+ proteins (21, 23, 39).

The NorQ protein alone has an ‘uncoupled’ (from target remodeling) ATPase activity of ~100 nmol mg^−1^min^−1^ (*k*_cat_ ~ 25 min^−1^ for the hexamer). This is similar to the homologous *A. ferrooxidans* AfCbbQ2 and *H. neapolitanus* csoCbbQ proteins (activity ~30 nmol mg^−1^min^−1^) (31, 33), which both have a sequence identity of ~50 % to NorQ (see alignment in Fig. S6) but higher than that observed for the CoxD (32) MoxR protein in the absence of its interaction (VWA-domain containing) partner. This activity was lost (~2 %) in the NorQ^WB^ variant, confirming the importance of the Walker B motif.

The NorQ^WB^ preparation was more homogenous than the wildtype NorQ, which led to better cryo-EM results. We could build a model for the NorQ^WB^ hexamer with a nominal resolution of 8 Å that is good agreement with the crystal structures of the homologous csoCbbQ (33) and AfCbbQ2 (25) (Fig. 5), where the latter resolved also a density for the helix 2 insert (H2I), a flexible loop (residue 78-93) at the convex side not resolved in the csoCbbQ crystal structure. Our NorQ structure has clear density in this region which is well modeled on the AfCbbQ2 structure (25). Because we determined the NorQ structure by cryo-EM, this indicates that the H2I region has a largely defined structure also in solution and not only in tightly packed crystals. Interestingly, NorQ hexamers that were built without applying symmetry constraints show the existence of asymmetric rings (Fig. 4). Asymmetric ring conformations are observed for nearly all hexameric AAA+ protein structures determined to date, where each protomer often assumes a different conformation related to the status of the ATP-binding pocket being either ATP or ADP bound, in an intermediate state of ATP hydrolysis or nucleotide free. It is common that one subunit of the ring is displaced, nucleotide-free and known as the “seam” protomer. Examples of such AAA+ proteins are ClpX, Vps4, LonA (40–42) or Hsp104 and ClpB for which opening of the ring is essential for function (43, 44). Asymmetric ring conformations were also observed for the MoxR AAA+ protein RavA (45).

### NorD forms a complex with the NorQ hexamer

Co-expressed NorQ co-purified with His-tagged NorD, showing that they interact (Fig. 3a).

Expression of NorD was dependent on co-expression of NorQ, possibly because binding to NorQ makes NorD soluble and/or prevents its degradation. Similar observations were made for heterologous expression of the CbbQO proteins involved in RuBisCO activation (31, 33).

The NorQD complex was obtained in significantly better yield and less aggregation prone with the NorQ^WB^ variant, presumably because of the loss of ATP hydrolysis activity. Our data (Fig. 3a and Fig. 6) shows that NorQ^WB^ forms primarily hexameric oligomers also within a NorQ^WB^D complex. BN-PAGE for NorQ^WB^D showed a predominant ~310 kDa band in the absence of ATP, which was verified to contain both NorQ and NorD (by mass-spec) and it is consistent with binding of one copy of NorD (70 kDa) to hexameric NorQ (6x 30 kDa). The cryo-EM data (Fig. 6a) shows a ‘comma’-like attachment to the NorQ^WB^ hexameric ring, consistent with this stoichiometry.

The VWA-domain of NorD is, as observed for other VWA domains (29), presumably involved in promoting metal ion-dependent protein-protein interactions via its conserved MIDAS motif (see Fig. 7a and conservation in other MoxR-associated VWA-containing proteins in Fig. S4). Since exchanging residues in this motif (T534V or D562N) of NorD still allowed co-purification of NorQD but lead to inactivation of *c*NOR in the context of the full operon (Table 2), the MIDAS motif is likely to be involved in binding to the target *c*NOR, but not to binding NorQ.

In our cryo-EM map (Fig. 8), we assume that it is the VWA domain of NorD that ‘plugs’ the center of the NorQ ring and we fitted the predicted structure of NorD (using the NorQD prediction, see Fig. 7c) into this density. This is done in analogy with the suggestion from data on the homologous CbbQ MoxR protein with its VWA-domain-containing partner protein CbbO (25, 31), and for other reasons detailed below. In the predicted structure of NorD as well as of all other MoxR-associated VWA domains predicted in this study there are ‘fingers’ in the VWA domain (see Fig. S8) that have not been observed in VWA domains experimentally (see e.g. PDB ID: 6FPY (46), but note that the available high-resolution structural data does not include any MoxR-associated VWA domains). We suggest that the insertion of the VWA ‘finger’ into the MoxR central pore is a general property also for other VWA domain-containing partner proteins to the MoxR AAA+ family members (e.g. CbbQ/O, RavA/ViaA), something that has to our knowledge not been observed or suggested before. We also suggest that this binding mode extends to eukaryotic AAA+ proteins in complex with VWA/MIDAS based on our predictions of the VWA domains of Rea1, ChlD and VWA8 (Fig. S14). In addition, we find it likely that the unassigned density (directly adjacent to a resolved VWA globular domain) observed in a cryo-EM structure of human Mdn1 (47) belongs to this VWA ‘finger’ (see Fig. S14c for the predicted fold of the Mdn homologue Rea 1 VWA domain). Binding of this region to the Mdn1 central pore was suggested previously, the ‘finger’ residues, however, are only partly resolved and resemble more an unstructured loop in the crystal structure (48). Interestingly, the ‘finger’ is missing in the well-studied VWA domains of e.g. integrins (see integrin-like ITI-HC1, Fig. S14e) and they also do not interact with AAA+ motors.

This binding mode would also explain mutagenesis results for CbbQ, where exchanging residues in the H2I region was found to lead to ‘uncoupling’ of the ATPase activity from the requirement for binding the CbbO partner (25), which we now can interpret in terms of the VWA ‘finger’ (of CbbO)/H2I (of CbbQ) interaction.

Binding of the VWA ‘finger’ to the center of the NorQ ring pore is also analogous to the McrC partner binding to the McrB AAA+ protein (26, 37), where the McrB hexamer ring structure shows high structural similarity to the NorQ/CbbQ MoxR proteins (24) although the sequence similarity is only ~20 %. The McrC partner protein (see Fig S14f) has no sequence homology to the VWA domain proteins. This binding mode presumably also rules out direct ‘threading’ of the target protein (*c*NOR) through the NorQ pore, as also suggested for McrBC (37).

From the cryo-EM NorQ^WB^D data (Fig. 6b), it is also clear that there is asymmetry in the NorQ hexamer and that this asymmetry has a defined relation to the binding of NorD, such that the ‘seam’ NorQ subunit appears to be the protomer least tightly bound to the NorD VWA. We suggest that this is related to the binding of the VWA domain ‘finger’ in the center of the NorQ ring, indicated also from the unassigned density (Fig. S11) at the convex (bottom) side of the NorQ ring that could be due to displacement of the NorQ H2I region in the protomer(s) tightly interacting with the VWA ‘finger’. This asymmetry would then be linked to the propagation of the ATPase motor function to the VWA domain and onto the target *c*NOR, as the ‘finger’ (at the bottom of the NorQ ring) is connected to the MIDAS residues (on the top) via a single β-strand (Fig. S15).

The N-terminal domain (unresolved in the final cryo-EM map) of NorD presumably sits on the side of the ring, as indicated by the ‘comma’ observed in the cryo-EM 2D classes (Fig. 6a) and the AlphaFold modelling (Fig. 8d, Fig. S9). We suggest its role is primarily to mediate specificity towards the *c*NOR target, supported by the observation that the N-terminal domain is not conserved and shows high variation in length in VWA-domain proteins associated with MoxR ATPases. Future higher-resolution structures resolving also the N-terminal domain of NorD should elucidate these issues.

### Mechanism of NorQD-catalyzed cNOR maturation

Intact NorQ AAA+ and NorD VWA domains and NorQ ATP hydrolysis capability (Table 1), are all required for assembly of active *c*NOR (Table 2). These pieces of evidence all support an ATP hydrolysis-driven motor function of NorQD in order to remodel *c*NOR to allow Fe insertion. An intact NorD VWA MIDAS motif was also essential for expression of active *c*NOR (Table 2), but not for binding to NorQ (see above). These observations are consistent with how we have modelled the cryo-EM density (Fig. 8) and indicate that NorD uses its VWA MIDAS motif to bind *c*NOR.

VWA domains often use their MIDAS-coordinated metal ion to coordinate to an acidic surface residue or ‘patch’ on the target protein (as the sixth metal ligand). We identified D220 and E222 on the NorB surface as good candidates for interacting with NorD, since their exchange yielded inactive *c*NOR, lacking Fe_B_ (Table 2). These effects are remarkable and highly unlikely to be caused by any direct effects on the Fe_B_ environment as both residues are positioned far from it (Fig. 9). Interestingly, the D220 and E222 residues are both located at the cytoplasmic end of TM helix VII which further up has two Fe_B_-coordinating histidines (H253/H254, see Fig. 9). Conformational changes in this helix induced by the NorQD motor function could then affect accessibility to and/or affinity for iron in the Fe_B_ binding site.

**Figure 9:**
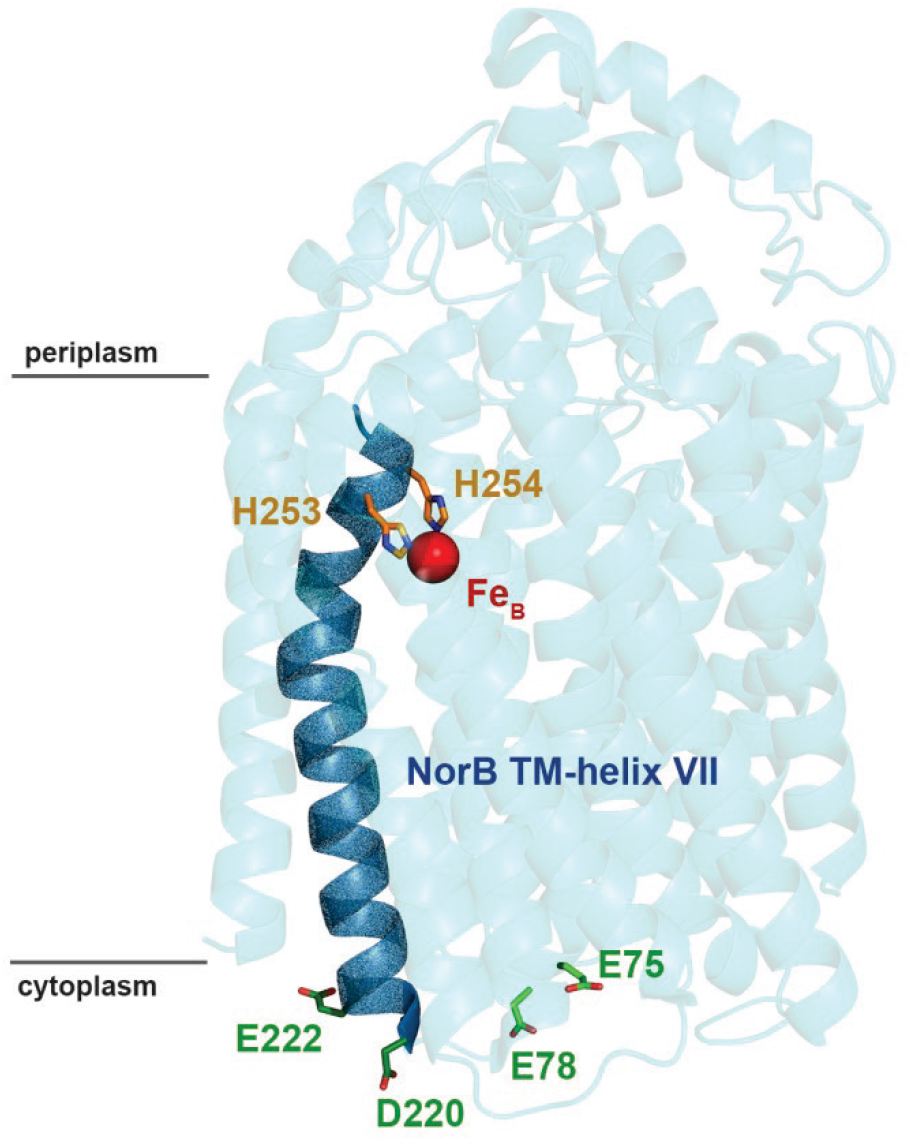
Mutational analysis of exposed acidic residues on the cytoplasmic surface of NorB. Residues investigated by mutagenesis (green) and Fe_B_-coordinating residues (orange) from TM helix VII (dark blue) are shown in stick representation. Picture based on Ps. aeruginosa cNOR (PDB: 3O0R, (5)) but amino acid numbering from the P. denitrificans cNOR sequence.

We found that the NorQD complex has lower ATP hydrolysis activity (Table 1) than the ‘uncoupled’ activity of NorQ alone. Thus basal NorQ activity is inhibited when in complex with the NorD, which is different from the situation in CbbQO, where the CbbQ ATPase activity is stimulated by the presence of CbbO (31). This difference is possibly exerted by the unrelated N-terminal domains of NorD/CbbO binding to the different final targets; for Rubisco the ‘after catalysis-inhibited protein’ and for *c*NOR presumably an assembly intermediate (see below).

Our modeled insertion of the NorD VWA domain ‘finger’ into the NorQ central pore explains in a straight-forward way how the ATP hydrolysis activity of NorQ is affected by the interaction in analogy with the influence (in their case stimulatory) the McrC binding was shown to have on the McrB hexamer (26, 37) where there is direct influence on the active nucleotide hydrolysis site by the insertion of the McrC ‘finger’. The NorD inhibition of the ‘uncoupled’ NorQ ATPase activity would then have to be relieved when NorD in turn binds to the ‘correct form’ of the *c*NOR target, since we know the ATPase function is necessary for *c*NOR activation (Table 2).

As the expression of *c*NOR in the absence of NorQD yields folded and hemylated *c*NOR lacking the Fe_B_ cofactor (2), we considered the possibility that NorQD could remodel the fully formed *c*NOR complex in order to open up a cavity that allows uptake and binding of Fe. Our re-activation experiments in crude cell extracts (Table S3) did lead to an increase in residual activity and non-heme Fe levels of Fe_B_-less *c*NOR in membranes, but only 2-2.5-fold. This result together with the lack of effect on the NorQD ATPase activity in the presence of purified Fe_B_-less *c*NOR suggests that the main target of the NorQD chaperone complex is a (temporarily formed) *c*NOR assembly intermediate and that they work inefficiently on the final *c*NOR product.

We previously showed that NorQD functions in non-heme Fe_B_ cofactor insertion into *c*NOR (2), but there was no biochemical data on the NorQD proteins. Based on the experimental data presented here, we can now suggest the following schematic mechanism for NorQD-catalyzed insertion of the Fe_B_ cofactor (Fig. 10): 1. The MIDAS motif of the NorD VWA domain binds to the surface residues D220/E222 on the TM helix VII of NorB in the apo-*c*NOR (without Fe_B_ bound). Presumably, also the N-terminal domain of NorD binds to *c*NOR in order to provide specificity and/or a ‘statoring’ function. 2. Fueled by ATP hydrolysis in NorQ, conformational changes in the ‘finger’ of VWA that extends through the central pore of the NorQ hexamer are propagated to the MIDAS surface residues and onto the *c*NOR surface opening/keeping open a channel/cavity around the TM helix VII that extends to the Fe_B_ binding site. Fe^2+^ (delivered from free Fe^2+^ in the cytosol or via iron chaperones/ferritin (49), but unlikely to involve the NorQD complex) can now access the Fe_B_ binding site. 3. When Fe_B_ is bound, conformational changes in the binding site could propagate back to the surface residues D220/E222 weakening the affinity for the MIDAS of the NorD VWA domain and the NorQD complex dissociates leaving the *c*NOR fully assembled and Fe_B_-bound.

**Figure 10:**
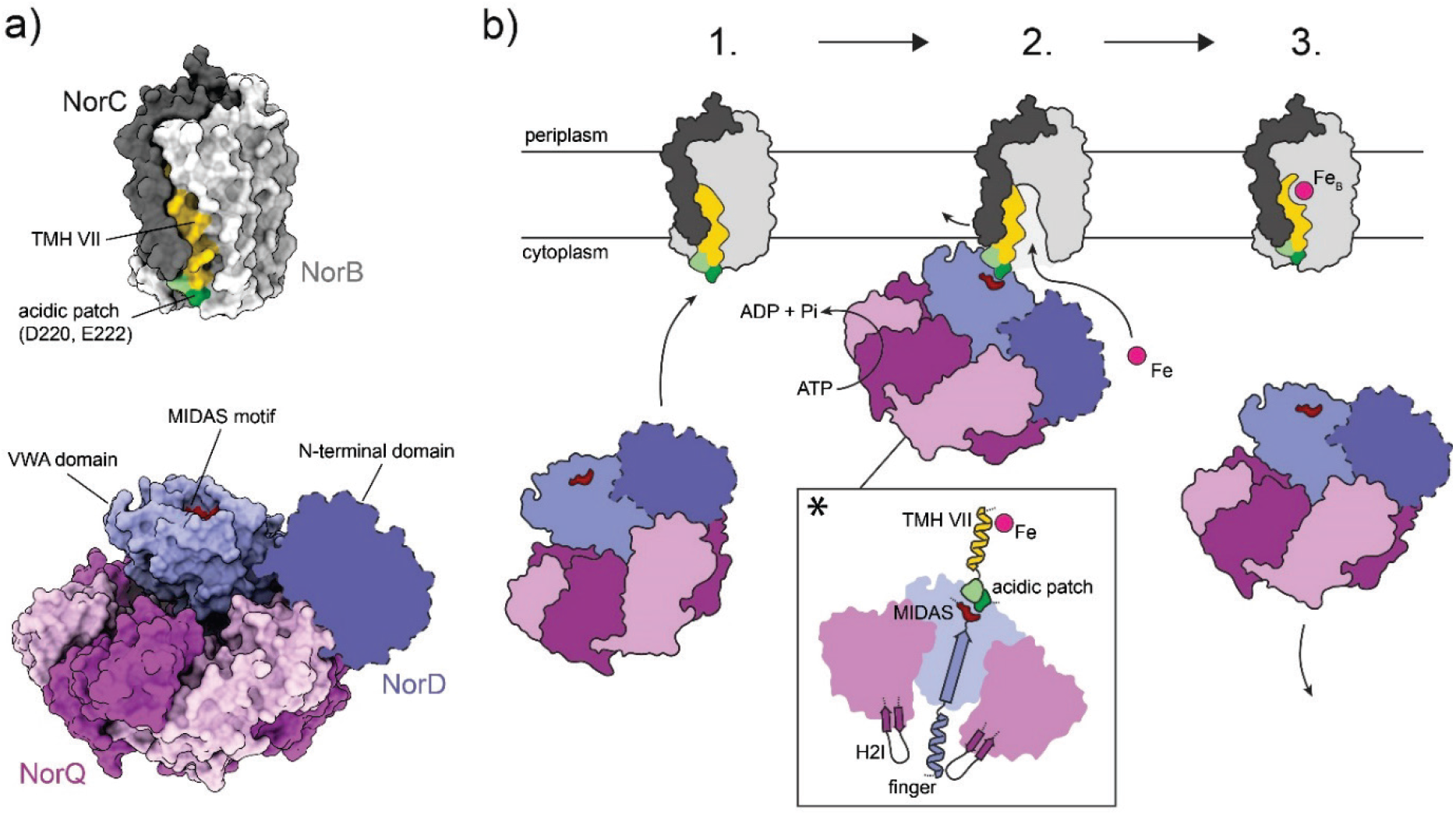
Proposed mechanism of cNOR activation by NorQD based on the presented data. a) Structure models for the P. denitrificans cNOR based on PDB ID: 3O0R (5), overall arrangement of the NorQD complex based on our cryo-EM data, detailed structures of NorD and NorQ protein based on 6L1Q (CbbQ for NorQ) and the predicted structure of NorD (Fig. 7a), respectively. b) Step 1: Hexameric NorQ (violet) uses one NorD (blue) as an adaptor protein in order to bind the apo-form of cNOR (grey, the suggested ‘immature’ nature of this form indicated by the larger accessibility of the acidic patch). Interaction of NorQD with cNOR occurs via the MIDAS site on the NorD VWA domain binding to D220/E222 on NorB (light grey). Step 2: NorQ induces ATP hydrolysis-powered conformational changes in cNOR via the ‘finger’ from the NorD VWA domain inserted into the hexamer pore. These conformational changes would then propagate from the NorD VWA via the NorB surface further up the TM helix VII, modulating accessibility and/or affinity of the Fe binding site. The inset (*) shows how the ‘finger’/H2I region in our model is structurally indirectly connected to the Fe_B_ site in cNOR. Step 3: Once cNOR has acquired the Fe_B_ cofactor and gained/restored functionality, the NorQD complex unbinds cNOR.

In summary, this study provides the first and novel insight into the specific mechanism of action of the NorQD-catalyzed insertion of the Fe_B_ into *c*NOR. In doing so, it also sheds light on the properties and mechanisms for the AAA+ MoxR-VWA protein interactions and mechanisms in the general context of this understudied class of enzymes.

## MATERIALS AND METHODS

### Cloning, expression and purification

*c*NOR was over-expressed in *Escherichia* (*E*.*) coli* from plasmids containing the complete *nor* operon (pNOREX-His) or in absence of the *nor* accessory genes (pNOREX-HisΔQDEF) (2, 50). For heterologous co-expression of NorQ and NorD an expression vector (pET-QhD) was constructed by cloning the *P. denitrificans norQ* and *norD* genes into a pET21a backbone. For both genes the native ribosome binding site (RBS) was exchanged for the T7 RBS (sequence: aaggag) and for NorD an N-terminal 6-His tag was introduced. We also expressed NorQ with an N-terminal 6-His tag in absence of NorD from a plasmid pET-hQ based on a pET21a backbone. The ‘ultrapure’ NorQD complexes were produced from a modified pETDuet-1 (Novagen) co-expression vector containing two multiple cloning sites (MCS). The original C-terminal S-tag located 3’ to the second MCS was exchanged for the Strep-tag II sequence. The NorD sequence was inserted into the first MCS and 3’ to the existing His-tag sequence and NorQ was cloned into the second MCS and 5’ to the Strep-tag II sequence via conventional restriction/ligation. This allowed for over- and co-expression of NorD with a N-terminal 6xHis tag and of NorQ with a C-terminal Strep-tag II. For site-directed mutagenesis a modified version of the QuickChange™ method and conventional cloning techniques were used. Correct assembly of pET-QhD and pET-hQ and site directed mutagenesis was confirmed by sequencing (Eurofins Genomics).

For *c*NOR expression pNOREX constructs were co-transformed together with pEC86 (encoding cytochrome *c* maturation factors(51)) into *E. coli* JM109 as described in (50). The growth procedure and His tag-based purification of *c*NOR is essentially described in (52, 53) with modifications of the purification protocol (2). For co- and over-expression of NorQ and NorD and individual expression of NorQ corresponding plasmids (pET-QhD respectively pET-hQ) were transformed into *E. coli* BL21 (DE3) and all growth media were supplemented with 100 mg/L ampicillin. 800 mL LB medium was inoculated with 8 mL of pre-culture and shaken at 180 rpm and 37 °C.

NorQ was grown at 30°C and shaken at 180 rpm overnight, without IPTG induction. NorQ^WB^ was induced with 1 mM IPTG and grown at 30°C and shaken at 180 rpm overnight. The cells were harvested by centrifugation and all the subsequent steps were performed on ice or at 4°C. NorQ cell pellets were suspended and diluted 4x in 100 mM HEPES, pH 7.6, 600 mM NaCl, 1 mM PMSF. NorQ^WB^ was suspended and diluted 4x in 100 mM HEPES, pH 7.6, 1 mM PMSF. Dilution was followed by cell disruption at 22 kpsi (Emulsiflex C3 Homogenizer) and centrifugation. 20 mM imidazole was added to the supernatant and the proteins were purified on a Hitrap HP column (5 mL GE/Cytiva), eluted with an imidazole gradient (50 mM to 500 mM) in 20 mM HEPES, pH 7.6, 300 mM NaCl, 5% (v/v) glycerol. The sample was dialyzed (12 kDa cut-off) against 20 mM HEPES, pH 7.6, 150 mM NaCl, 5% (v/v) glycerol. Concentration was determined using A280.

NorQD was grown at 25°C and shaken at 180 rpm overnight, without IPTG induction. NorQ^WB^D was induced with 0.1 mM IPTG, grown at 16°C and shaken at 180 rpm for 40 h. The purification proceeded as for NorQ and NorQ^WB^ with some changes: after harvest, the cells were diluted in 4x 50 mM TRIS, pH 8, 50 mM NaCl, supplemented with one pill cOmplete, EDTA-free Protease Inhibitor Cocktail. The purification buffer contained 20 mM TRIS/HCl, pH 8, 300 mM NaCl, 10% (v/v) glycerol and imidazole concentration ranging from 50 to 500 mM. The sample was dialyzed against 10 mM TRIS, pH 8, 150 mM NaCl, 10% (v/v) glycerol.

Double-tagged (Q-Strep, D-His) NorQD and NorQ^WB^D were grown and purified as described above with an additional purification step on Strep-Tactin superflow column (iba). The sample was eluted with 2.5 mM desthiobiotin, followed by His-tag purification as before.

For SDS-PAGE, protein samples were diluted in NuPAGE™ LDS sample buffer (Invitrogen) After incubation for 10 min at 95 °C, the sample was loaded and run on NuPAGE™ 4-12% Bis-Tris gels (constant 100 V, room temperature) and stained with Coomassie Brilliant Blue or silver staining. Identification of individual protein bands was supported by mass spectrometry analysis performed by the SciLifeLab Mass Spectrometry Facility at Uppsala University using nano-LC-MS/MS.

Blue native (BN) PAGE was performed with pre-cast NativePAGE™ 4-16% Bis-Tris (Invitrogen) gels according to the manufacturer’s instruction. The gels were run at 4 °C in two stages: 1) 60 min at 150 V, 2) 45 min at 250 V. The gels were stained with Coomassie Brilliant Blue.

NorQ^WB^ for Cryo-EM was produced as described above, with some exceptions. The cells were harvested 6 h after induction and diluted 4x in 100 mM HEPES, pH 7.6, 20 mM β-mercaptoethanol. After cell disruption and centrifugation, 20 mM imidazole and 10% (v/v) Ni-NTA agarose resin was added to the supernatant and the mixture was incubated under rotation for 2 h. The resin was added to a gravity-controlled column and washed with 5 CV (column volumes) of purification buffer (20 mM HEPES, pH 7.6, 300 mM NaCl, 5 % (v/v) glycerol, 20 mM β-mercaptoethanol) containing 20 mM imidazole. The protein was eluted with 2 CV of purification buffer containing 250 mM imidazole. The sample was dialyzed (12 kDa cut-off) against 20 mM HEPES, pH 7.6, 150 mM NaCl, 2 mM DTT overnight. Finally, all protein samples were concentrated with centrifugal filters (10 kDa cut-off) and concentrations were determined using the Pierce™ BCA Protein Assay Kit.

Purified protein samples were analyzed with size exclusion chromatography (SEC). Approximately 500 μg protein was run on a Superdex 200 10/300 GL-column (Cytiva) at 0.3 mL/min in 20 mM TRIS/HCl pH 8, 300 mM NaCl, 10% (v/v) glycerol. Detergent solubilized membranes were prepared by using the same *c*NOR expression system described above. Mutants of NorB, NorQ or NorD were produced by genetic modification of pNOREX-His. After induction with 1 mM IPTG, the cells were grown in 25 mL TB (containing 100 mg/L ampicillin, 30 mg/L chloramphenicol) overnight at 30 °C, 180 rpm. Cells were pelleted by centrifugation and suspended in 1 mL of 100 mM TRIS/HCl, pH 7.5, 50 mM NaCl, 1 mM EDTA. The cells were sonicated for 2 min (30s on/30s off cycle) in an ice bath followed by addition of 1 % DDM. Insoluble cell debris was removed by centrifugation (20,000 g, 10 min) and *c*NOR concentrations in the supernatant were determined spectrophotometrically (ɛ_550 reduced-oxidized_ = 28 mM^−1^ cm^−1^). *c*NOR is the only enzyme with appreciable NO-reduction activity as well as the only *c*-heme containing protein in the membranes, making normalization to *c*NOR content straight-forward.

### ATPase activity measurements

ATPase activity was measured using the pyruvate kinase coupled assay (54). The reaction mixture contained 50 mM TRIS/HCl, pH 8.0, 150 mM NaCl, 15 mM MgCl_2_, 2.5 mM ATP, 1 mM phosphoenolpyruvate, 0.3 mM NADH, 12 U/mL pyruvate kinase, 12 U/mL lactate dehydrogenase. After addition of 0.15 mg/mL protein the ATPase activity was measured by following NADH consumption photometrically (ε_340_ = 6.22 mM^−1^ cm^−1^) at 22 °C. Because of the tendency of the NorQ and NorQD complex to aggregate at higher concentrations, the sample was centrifuged, the protein concentration determined right before the activity measurements and the stock solutions kept at 0.5-1mg/ml or lower.

In order to avoid potential artifacts in the optical assay from competing aggregation reactions of the samples, ATPase activity was also measured using the malachite green end-point assay (55). The initial reaction mixture contained 50 mM TRIS/HCl, pH 8, 150 mM NaCl, 10% (v/v) glycerol, 20 mM MgCl_2_, 2 mM ATP. 1 min after addition of 0.03 or 0.05 mg/mL protein, 109 μL color reagent (0.034% malachite green oxalate, 1% ammonium molybdate tetrahydrate, 0.04% Triton-X) was added to 27.3 μL of the initial reaction mixture, and the sample was vortexed. After 1 min, 13.5 μL 34% citrate solution was added and the sample was vortexed. After 5 minutes, the absorption at 620 nm was measured and the phosphate concentration determined using a standard curve. The experiment was repeated at regular time points after addition of protein, and the ATPase activity determined as the increase of phosphate concentration over time. Measurements were performed at least two different protein concentrations in order to avoid artefacts from aggregation reactions, the rate of which should increase with protein concentrations.

### NO multiple turnover measurements

NO reduction activity was measured for detergent solubilized membranes and for purified *c*NOR (see above). NO reduction measurements were performed as described in (56) with the following modifications: 25 nM *c*NOR was added already before deoxygenating the buffer solution. The reaction was started by addition of 10 μM horse heart cyt. *c*, 0.5 mM TMPD and 3 mM ascorbate after injection of NO. The measurements were performed at 22 °C.

### Re-activation experiments

The *nor* operon proteins were expressed as described above from pNOREX-His and pNOREX-HisΔQDEF plasmids. After cell lysis and subsequent centrifugation, the membrane fraction and soluble fraction was saved. In order to assess a possible re-activation effect of soluble NorQ and NorD proteins on membrane bound, Fe_B_-lacking *c*NOR, the membrane fraction and supernatant were re-united according to the scheme in Table 2. The samples were homogenized and incubated for 1.5 h at room temperature followed by sedimentation of the membrane fraction by centrifugation. *c*NOR was then purified from membranes as described above. Protein concentrations were determined spectrophotometrically (ɛ_410_ = 311 mM^−1^ cm^−1^ (57)) followed by NOR activity measurements and non-heme iron determinations.

### Non-heme iron determinations

The non-heme iron content was measured using the ferene method (58, 59). To 60 μL protein solution (8 μM) 6 μL of 37 % HCl were added. The sample was incubated for 5 min and precipitated protein was removed by centrifugation (20,000 g, 15 min), followed by addition of 100 μL of 3 M sodium acetate and 10 μL of 1 M ascorbic acid (pH 5-6). 10 μL of 3 mM ferene was added and after 5 minutes, the non-heme Fe concentration was measured photometrically (ɛ_593_ = 35.5 mM^−1^ cm^−1^). The signal was corrected for the background of contaminating Fe dissolved in the buffer as well as for background absorbance in the sample before ferene addition.

### Cryo-EM

For NorQ alone, samples contained 1 mg/mL protein in 20 mM HEPES, pH 7.6, 150 mM NaCl, 2 mM DTT. 3.5 μL of the sample was blotted on CF-2/2-3Cu-T-50 grids (glow discharge for 40 s, blotting for 3 s, blotting chamber at 4 °C, 100 % humidity).

For co-purified NorQ^WB^D, cryo-EM samples contained 1 mg/mL protein in 50 mM TRIS/HCl, pH 8.0, 150 mM NaCl, 10% glycerol.

The dataset of NorQ without NorD was collected on a Talos Arctica (Thermo Fisher Scientific) using a Falcon3 detector (Thermo Fisher Scientific) in counting mode. The exposure time was 80 s and the movie frames 42 with 1.24 e^−^/Å^2^ per frame. The data were collected at a magnification of 150k giving a pixel size of 0.99 Å. On-the-fly data processing was performed as for the NorQD complex and further image processing using Relion 3 (60) integrated into the Scipion frame.

Around 130,000 particles were picked automatically on-the-fly using the Xmipp pattern matching approach protocol (61). Upon 2D classification, particles belonging to good 2D classes both representing end and side views were selected in a dataset of around 50,000 particles. Two 3D initial models were generated using the *ab initio* stochastic gradient descent method implemented in Relion3. The two models looked virtually identical and one of the two was used as reference for 3D classification into two classes. The 3D classification allowed to further clean the dataset and 90% of the particles contributed to a good 3D class while the rest segregated into a ‘junk’ 3D. The 42,000 particles belonging to the good 3D class were refined using C6 symmetry into an 8Å map and, with no-symmetry applied, into a 11Å map. The C6 map was used for 3D classification while relaxing the symmetry.

Data acquisition of NorQD was performed on a Titan Krios G3i (Thermo Fisher Scientific) microscope using a K3 detector (Gatan, Ametek) in super-resolution hard-binned mode, using a dose rate of 12 e^−^/px/sec and a pixel size of 0.8464 Å. The total dose was 40 e^−^ and the movie frames 40. On-the-fly data processing was performed using the cryoSPARC Live suite (62) and subsequent image processing was continued in cryoSPARCv 3.0. From 23018 micrographs, ~14 million particles were blob-picked on-the-fly, but only ~500.000 were selected using 2D classification and three ab-initio models were created with and without C6 symmetry applied. The asymmetric models all contained extra density inserted in the NorQ ring and a lateral “coma-like” protrusion later assigned to NorD. The dataset was subjected to extensive 3D classification and using the six-fold symmetric model helped in angles assignment. This was due to substantial preferential orientation of the available views so that an isotropic map was only obtainable to a nominal resolution of 5Å using non-uniform refinement in cryoSPARC. By inspection, the map is more consistent with a 10Å resolution where AAA+ subdomains are easily recognisable as well as the VWA domain. While the ‘comma’ density was present in the ab-initio models, it was lost in the further refinement. Based on other evidence we interpret the ‘comma’ as the N-terminus of NorD and it is most likely flexible. More isotropic data is required to be able to determine a high-resolution structure and to distinguish different conformations.

### Prediction of single-chain NorD and NorQD Complexes using AlphaFold

The AlphaFold prediction (34) of the structure of NorD alone (and related proteins, Fig. S7 and Fig. S8) was made using the web resource (63) at https://colab.research.google.com/github/deepmind/alphafold/blob/main/notebooks/AlphaFold.ipynb

All models of multimeric complexes shown in Results were built using AlphaFold-multimer original or updated version (35) with default settings. Here four multiple sequence alignments (MSAs) are created by searching Uniref90 v.2020_01 (64), Uniprot v.2021_04 (65) and MGnify v.2018_12 (66) with jackHMMER (67) and using Hhblits (68) to search the Big Fantastic Database (69) and uniclust30_2018_08 (70). Only proteins from UniProt are paired by using species information as described in Evans et al. (35). All of the created MSAs (one paired and three block-diagonalized) are used to predict the structure of a protein complex. Models are evaluated using the predicted template/model (pTM) score (71).

We also used the FoldDock protocol (72) to build complexes, however, for the resulting models only one out of five had the NorD in the center of the hexamer, and was thus considered not consistent with experimental data.

## Abbreviations

*c*NOR: cytochrome *c*-dependent nitric oxide reductase
MIDAS: metal ion-dependent adhesion site
VWA: von Willebrand factor Type A
qNOR: quinol-dependent nitric oxide reductase
HCuO: heme-copper oxidase
TM: trans-membrane (helix)
WT: wildtype *c*NOR
TMPD: N,N,N′,N′-Tetramethyl-p-phenylene-diamine
RBS: ribosome binding site
DDM: n-Dodecyl β-D-maltoside
cyt. *c*: cytochrome *c*
WB: Walker B (motif).

## Data availability

The cryo-EM density maps will be deposited in the Electron Microscopy Data Bank under accession codes EMD-XXX and EMD-YYY. All other datasets generated during and/or analysed during the current study are either included in the manuscript or supporting information file or available from the corresponding authors on reasonable request.

## ACKNOWLEDGEMENTS

This work was supported by grants from the Swedish Research Council (VR-2015-04512 and 2019-04124) to PÄ. The Cryo-EM data was collected at the Cryo-EM Swedish National Facility funded by the Knut and Alice Wallenberg, Family Erling Persson and Kempe Foundations, SciLifeLab, Stockholm University. The AlphaFold multimer computations were supported by the Swedish Research Council for Natural Science (VR-2016-06301 to AE), the Swedish E-science Research Centre, and the Knut and Alice Wallenberg foundation and the computational resources are supported by the Swedish National Infrastructure for Computing, grants: SNIC 2021/5-297, SNIC 2021/6-197 and Berzelius-2021-29. We are grateful to Patrick Bryant och Gabriele Pozzati for help with implementing the use of AlphaFold-multimer. The mass spectrometry analysis was supported by the Science for Life Laboratory Mass Spectrometry Based Proteomics Facility in Uppsala. The pET-Duet vectors for double-tagged NorQD were constructed in the lab of Petra Wendler (University of Potsdam).

## Author contributions

MK and PÄ conceived the study. MK, SA, MC and PÄ designed the experiments. MK, SA, and MC performed the experiments. MK, SA, MC and PÄ analyzed the data. AE performed the AlphaFold-multimer predictions and evaluated the models. MK and PÄ drafted the manuscript with parts contributed by SA, MC and AE. All authors revised and commented on the draft and on the final manuscript.

## Competing interests

The authors declare no competing interests.

## Notes

### Competing Interest Statement

The authors have declared no competing interest.

