## SupplementalInformation for "Insights into the structure-function relationship of the NorQ/NorD chaperones from *Paracoccus denitrificans* reveal shared principles of interacting MoxR AAA+/VWA domain proteins"

[%]Authors contributed equally

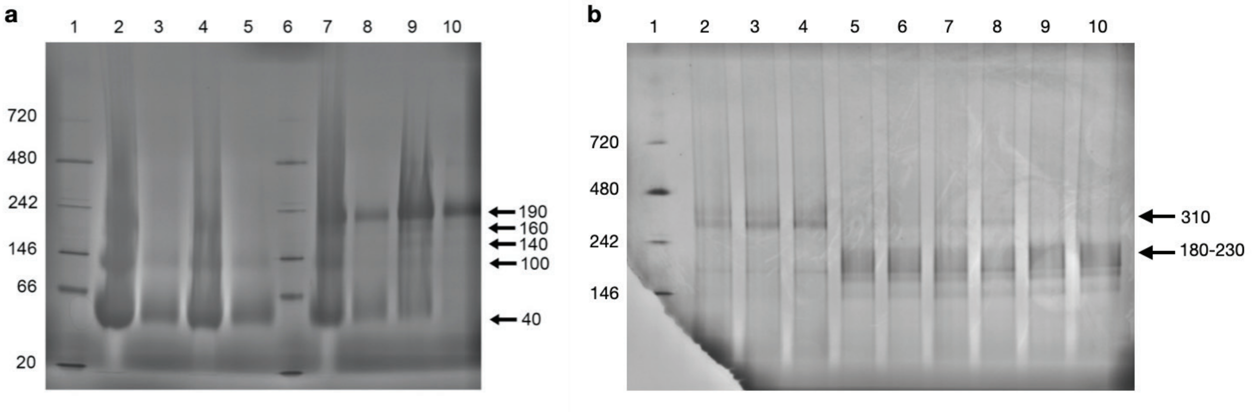


***Figure S1: Blue Native (BN-) PAGE gels.*** *Run at 4° C for 1 h at 150 V, followed by 30 min at 250 V. a) Blue Native (BN-) PAGE of wild type NorQ and NorQ$^{WB}$ with and without addition of ATP and $MgCl_2$. Original gel to Figure 1b in main text. Lane 1: Native Marker, 5 µL. Lane 2: NorQ, 29 µg. Lane 3: NorQ, 2.9 µg. Lane 4: NorQ, 14 µg, with 2 mM ATP and 20 mM $MgCl_2$, Lane 5: NorQ, 2.9 µg, with 2 mM ATP and 20 mM $MgCl_2$. Lane 6: Native Marker, 5 uL. Lane 7: NorQ$^{WB}$, 26 µg. Lane 8: NorQ$^{WB}$, 3.6 µg, Lane 9: NorQ$^{WB}$, 18 µg with 2 mM ATP and 20 mM $MgCl_2$. Lane 10: NorQ$^{WB}$, 3.6 µg with 2 mM ATP and 20 mM $MgCl_2$. Gel run for 1h at 150 V and 45 min at 250 V. b) Blue Native (BN-) PAGE of NorQ$^{WB}$D with different additions of ATP and $MgCl_2$. Original gel to Figure 2b in main text. Lane 1: Native Marker, 5 µL. Lane 2-4: NorQ$^{WB}$D, 10 µg, with 20 mM $MgCl_2$. Lane 5-6: NorQ$^{WB}$D, 10 µg, with 0.5 mM ATP and 20 mM $MgCl_2$. Lane 7-8: NorQ$^{WB}$D, 10 µg, with 1 mM ATP and 20 mM $MgCl_2$. Lane 9-10: NorQ$^{WB}$D, 10 µg, with 2 mM ATP and 20 mM $MgCl_2$.*

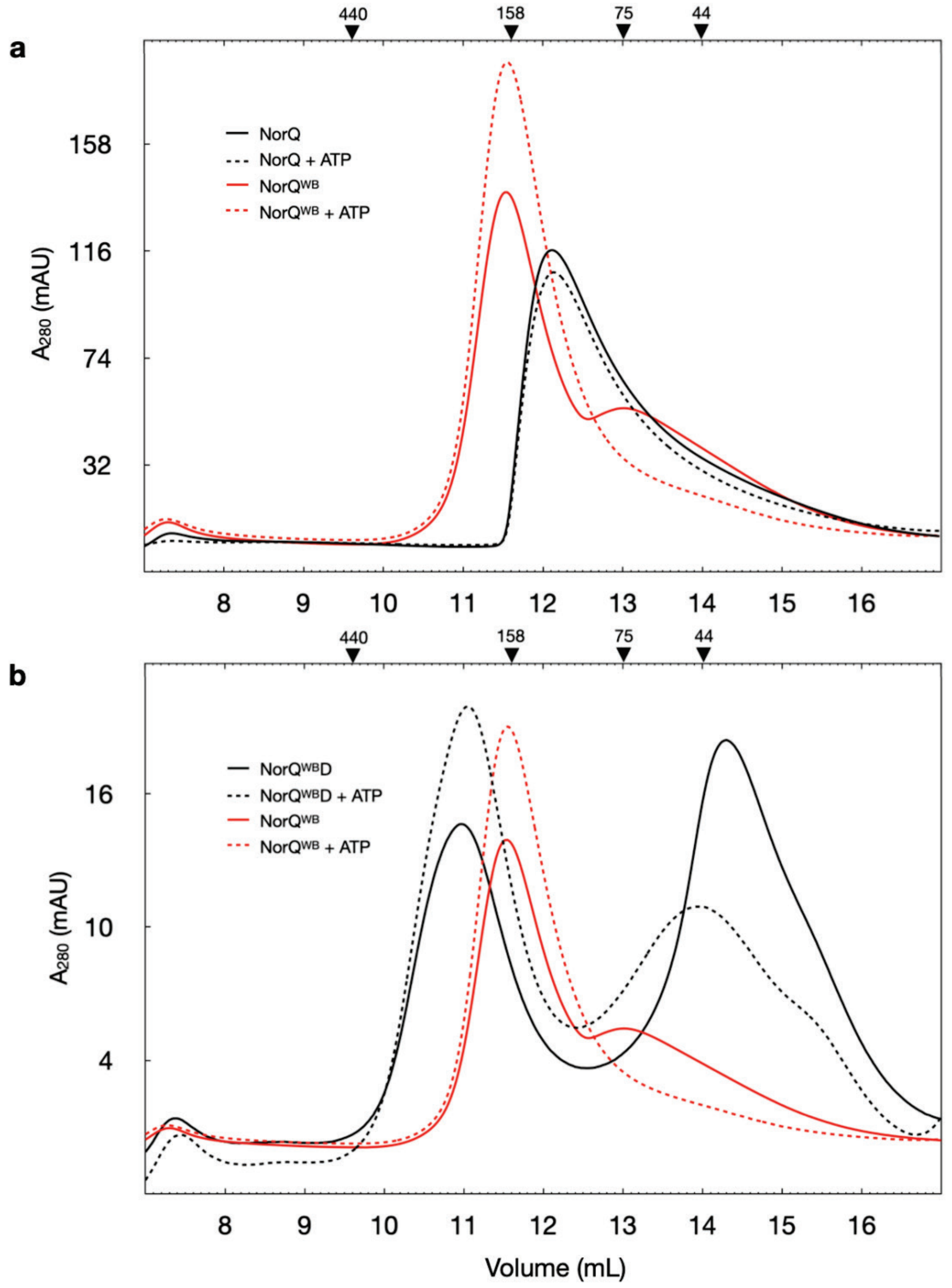


***Figure S2: SEC profiles.*** *a) NorQ (black solid) and NorQ with 2 mM ATP (black dashed). Both show one major peak, eluting at ~120 kDa. NorQ$^{WB}$ (red solid) showing two major peaks, eluting at ~160 kDa and ~70 kDa. NorQ$^{WB}$ with 2 mM ATP (red dashed) showing only the ~160 kDa peak. b) NorQ$^{WB}$D (black solid) showing two major peaks, eluting at ~210 kDa and ~40 kDa and NorQ$^{WB}$D with 2 mM ATP (black dashed), also two major peaks at ~210 kDa and ~50 kDa. The same NorQ$^{WB}$ profiles as in a) are shown for comparison. All samples were run on Superdex 200 10/300 GL and the markings on top are from a calibration with GE Gel Filtration HMW Calibration Kit.*

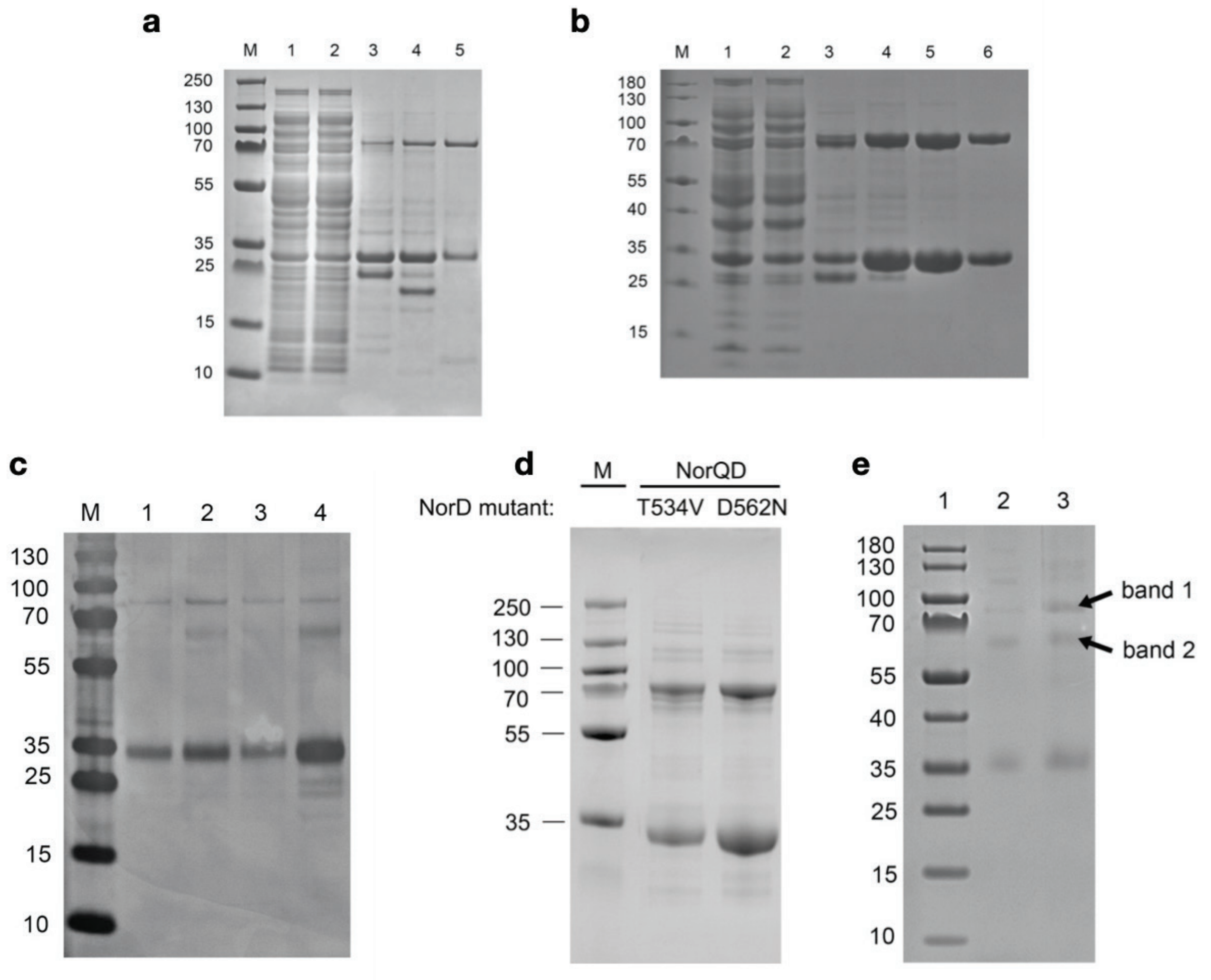


***Figure S3: Original SDS-PAGE gels.*** *All gels run in room temperature at 100 V. a) SDS-PAGE of NorQD purification. Original gel to part of Figure 2a in main text. Lane M: PageRuler Plus Prestained, 5 µL. Lane 1: Cytosolic suspension. Lane 2: Flow-through from Ni-NTA. Lanes 3-5; fractions from imidazole stepwise elution: 3: 150 mM imidazole, first peak. Lane 4: 150 mM imidazole, second peak. Lane 5: 200 mM imidazole. b) SDS-PAGE of NorQ$^{WB}$D purification. Original gel to part of Figure 2a in main text. Lane M: PageRuler Prestained. Lane 1: Cytosolic suspension. Lane 2: Flow-through from Ni-NTA. Lanes 3-6: Fractions from the imidazole gradient: Lane 3: 100-160 mM imidazole. Lane 4: 160-210 mM imidazole. Lane 5: 210-260 mM imidazole. Lane 6: 260-360 mM imidazole. c) SDS-PAGE on BN-PAGE bands from figure S1b. Original gel to Figure 2c in main text. Lane M: PageRuler Plus Prestained. Lane 1: Upper 310 kDa band from lanes 2-4 in S1b. Lane 2: Lower 310 kDa band from lanes 2-4 in S1b. Lane 3: 180-230 kDa band from lanes 2-4 in S1b. Lane 4: 180-230 kDa band from lanes 5-10 in S1b. d) SDS PAGE of co-purified NorQD after mutagenesis of the conserved MIDAS motif. Lane 1: PageRuler Plus Prestained. Lane 2: NorQD$^{T534V}$, 1 µg. Lane 3: NorQD$^{D562N}$, 1 µg. e) SDS-PAGE for mass spectrometry analysis of NorQD. Lane 1: Marker; Lane 2: not relevant; Lane 3: 290 kDa complex isolated from BN-PAGE. The gel bands analyzed by mass-spec are indicated by an arrow. The corresponding mass-spec data is shown in Table S1.*

***Table S1: Mass spectrometry analysis of selected bands from SDS-PAGE (see Figure S3e).***

| sample | detected protein | Nr. of identified peptides | coverage (%) | protein score |
|---|---|---|---|---|
| **band 1** | *Major components:* | | | |
| | NorD | 37 | 54 | 275 |
| | NorQ | 20 | 60 | 81 |
| | *E. coli contaminants:* | | | |
| | Histone family protein DNA-binding protein | 2 | 28 | 2 |
| **band 2** | *Major components:* | | | |
| | NorQ | 21 | 70 | 290 |
| | NorD | 32 | 53 | 84 |

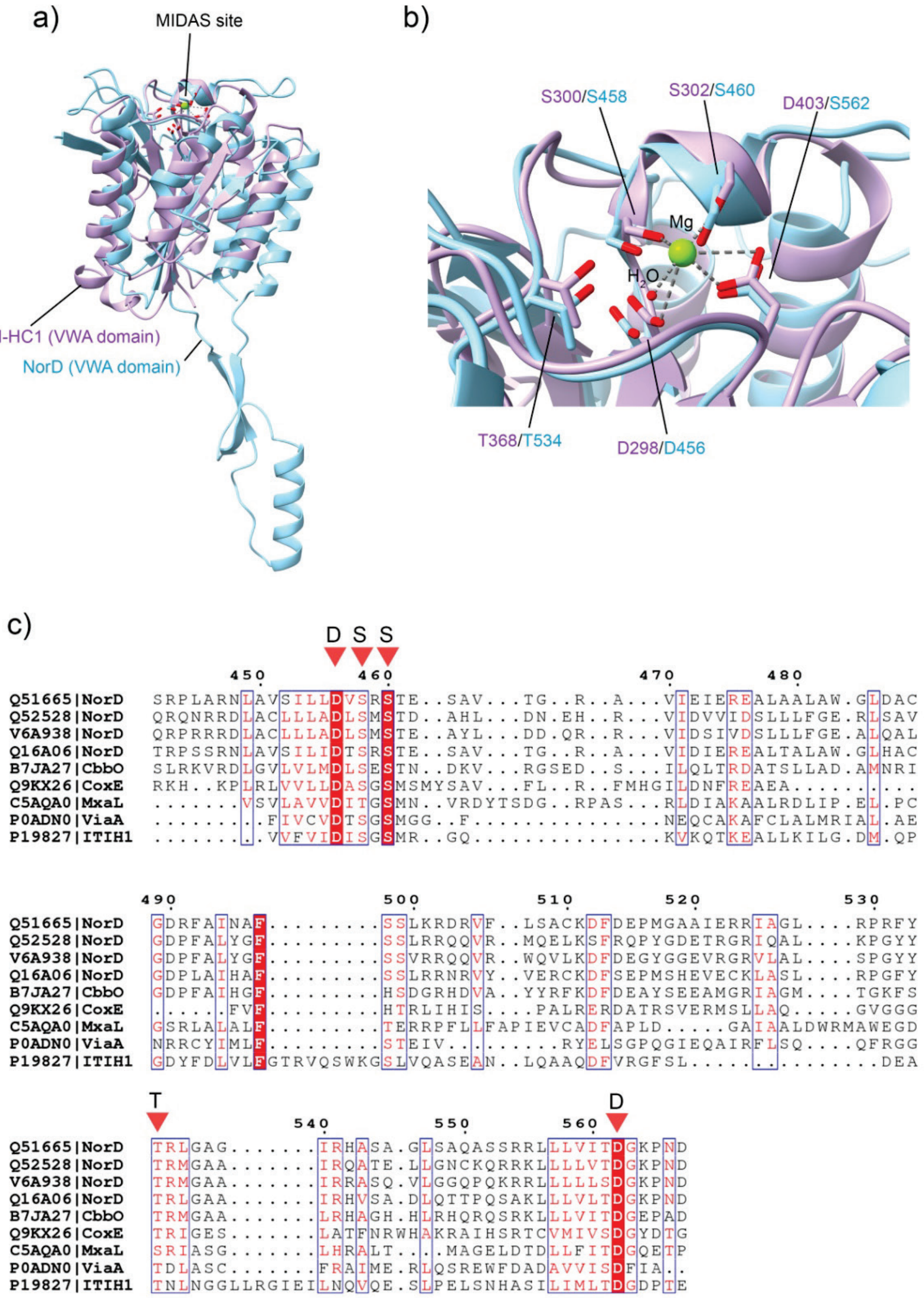


***Figure S4: Comparison of VWA domains and the MIDAS region from different organisms.*** *a) Predicted AlphaFold VWA domain model of NorD (cyan, Uniprot (UP): Q51665, see also Fig. 7a in main text) superimposed on the crystal structure of the Inter-alpha-inhibitor heavy chain 1 (purple, ITI HC1, PDB ID: 6FPY (1)). b) Enlargment of the conserved MIDAS region, with the residues shown to (for ITI, purple) or predicted to (for NorD, cyan) bind Mg (green) marked. c) Sequence alignment of selected VWA domain MIDAS regions: NorD from P. denitrificans (UP: Q51665), Ps. stutzeri (UP: Q52528), Ps. aeruginosa (UP: V6A938), and Roseobacter denitrificans (UP: Q16A06), CbbO1 from Acidithiobacillus ferrooxidans (UP: B7JA27), CoxE from Afipia carboxidovorans (UP: Q9KX26), MxaL from Methylorubrum extorquens (UP: C5AQA0), ViaA from E. coli (UP: P0ADN0) and human ITI-HC1 (UP: P19827). Residue numbers according to P. denitrificans NorD. The sequence alignment was made using T-Coffee (2, 3) and ESPpript 3.0 (4).*

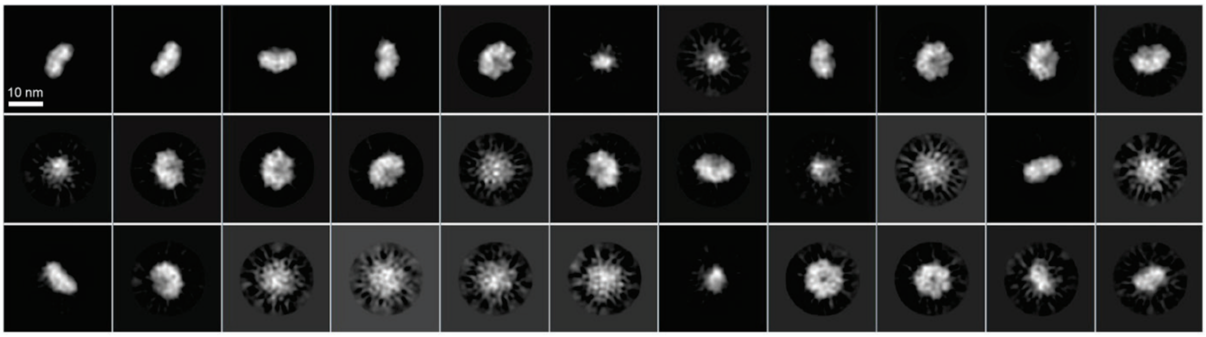


***Figure S5: 2D class averages of NorQ in cryo-EM experiments.*** *The 2D classes represent 50,000 particles out of a total number of 130,000 particles.*

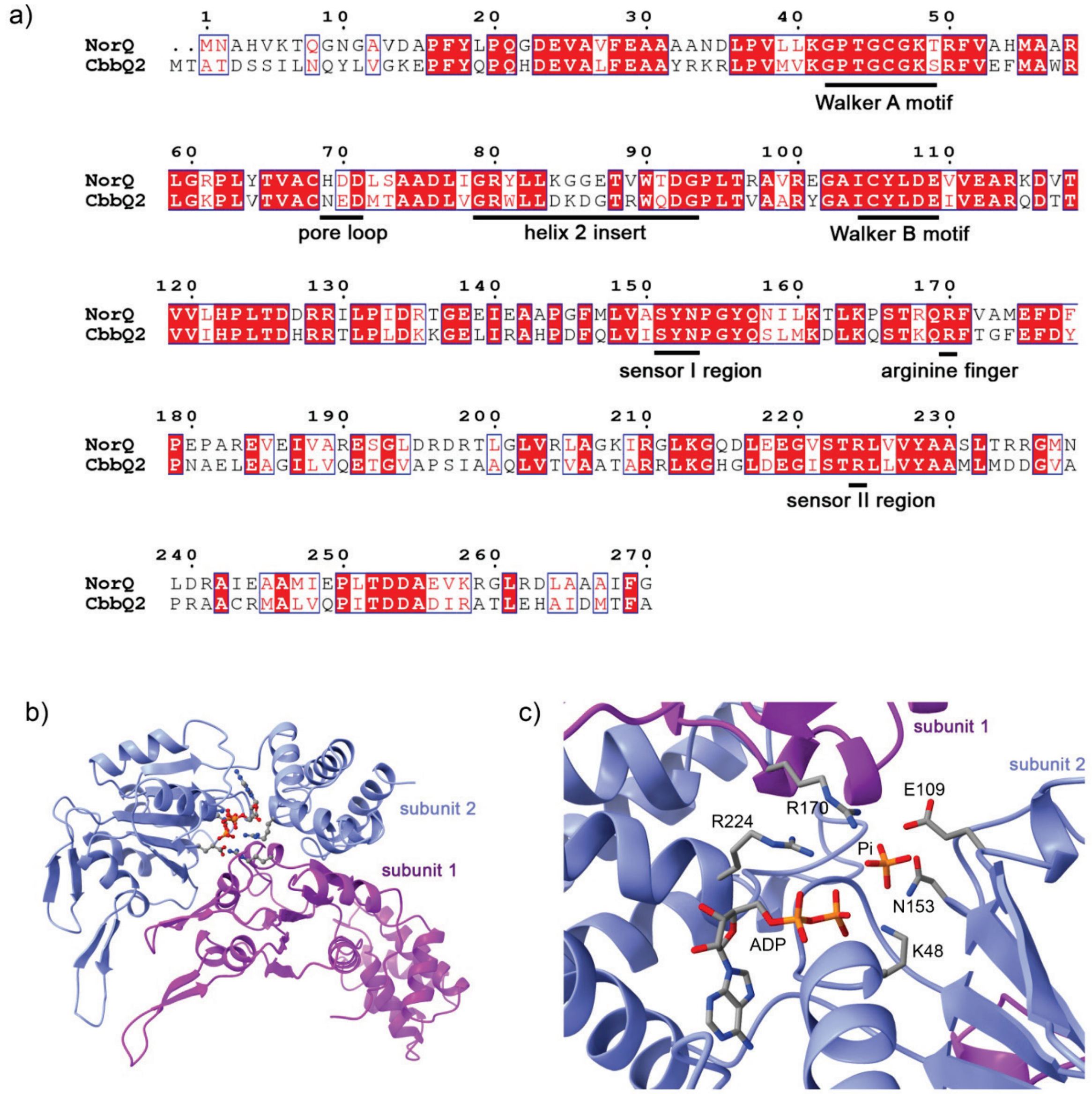


***Figure S6: a) Sequence alignment of AfCbbQ2 from A. ferrooxidans (Uniprot: B7J5E4) with NorQ from P. denitrificans (Uniprot: Q51664)****. The conserved features of the AAA+ domain are marked. The sequence alignment was made using Clustal Omega (3) and ESPpript 3.0 (4). b) Interaction of two adjacent NorQ subunits and formation of the active site (nucleotide and involved residues are shown in ball-stick representation). A homology model (SWISS-MODEL, (5)) of the P. denitrificans NorQ is shown based on the crystal structure of AfCbbQ from A. ferrooxidans (PDB ID: 6L1Q). The coordinates of the nucleotide and phosphate are taken from the AfCbbQ crystal structure after superimposition with the NorQ model. c) Close-up of the nucleotide binding pocket of the NorQ model formed between neighboring subunits.*

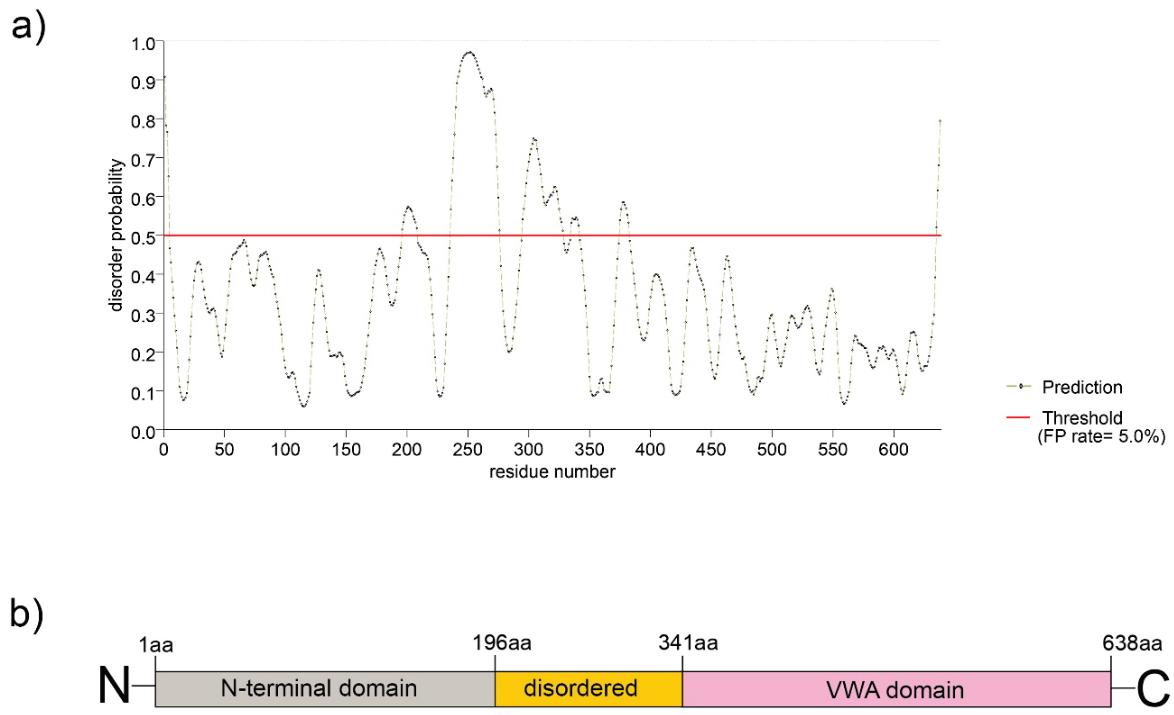


***Figure S7: Prediction of disordered regions and overall architecture for full length NorD from P. denitrificans.*** *a) Disordered regions predicted using the PrDOS server (6). b) Overview of the architecture of the NorD protein including the N-terminal domain (grey), disordered linker (yellow) and the C-terminal VWA-domain (pink) as predicted by AlphaFold (see Fig. 7a in main text).*

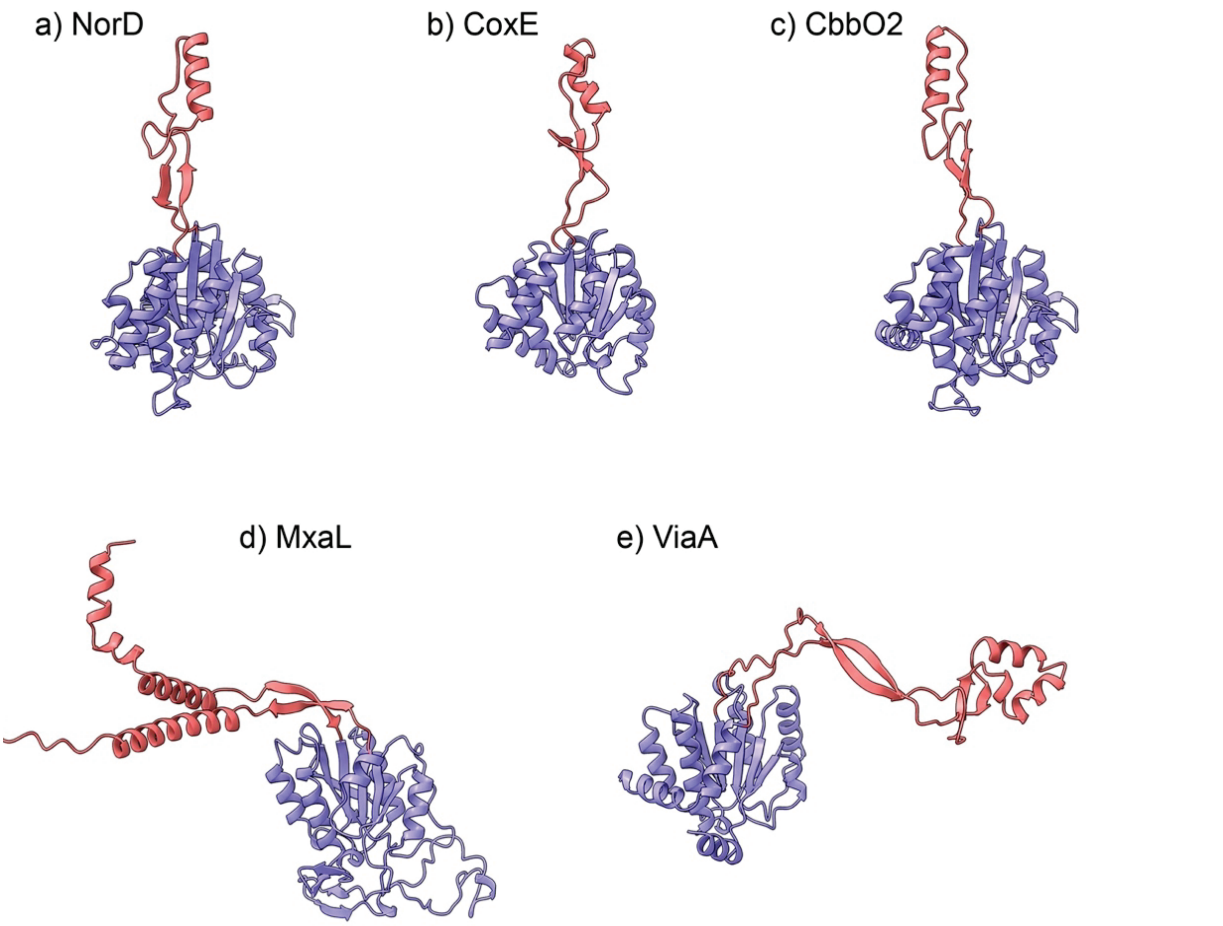


| *protein (VWA domain only)* | *Uniprot ID* | *residues VWA* | *sequence identity (%)* | *sequence similarity (%)* |
|---|---|---|---|---|
| ***NorD*** | *Q51665* | ***337-638*** | ***100*** | ***100*** |
| ***CoxE*** | *Q9KX26* | ***163-399*** | ***18*** | ***28*** |
| ***CbbO2*** | *B7J5E5* | ***463-759*** | ***31*** | ***51*** |
| ***MxaL*** | *C5AQA0* | ***1-336*** | ***12*** | ***21*** |
| ***ViaA*** | *P0ADN0* | ***195-483*** | ***18*** | ***30*** |

***Figure S8**: **Finger-like protrusions predicted for several MoxR-associated VWA domains and pairwise alignments.*** *The predicted VWA domains are all models produced by AlphaFold. VWA domains in blue and finger-like protrusions in red. a) NorD from P. denitrificans (Uniprot: Q51665), b) CoxE from Afipia carboxidovorans (Uniprot: Q9KX26), c) CbbO2 from Acidithiobacillus ferrooxidans (B7J5E5), d) MxaL from Methylorubrum extorquens (Uniprot: C5AQA0), e) ViaA from E. coli (Uniprot: P0ADN0). The table shows the results of pairwise alignments of the NorD VWA domain to the other VWA domains depicted in this figure. The models shown correspond to the sequence fragments used for the alignments. For the alignments EMBOSS Needle (7) was used.*

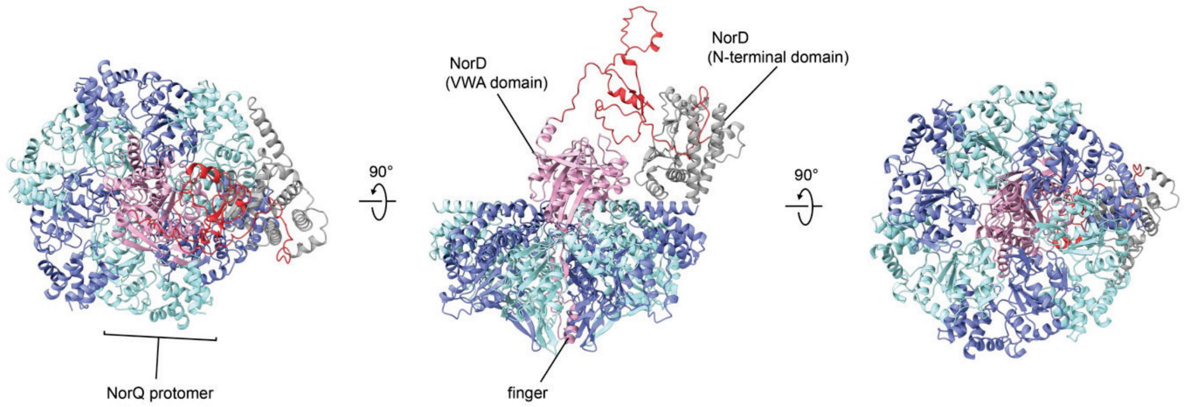


***Figure S9**: **AlphaFold multimer model of the full length NorD (one chain) and NorQ (six chains).*** *The NorQ protomers are colored in cyan and blue. The NorD is shown with the N-terminal domain in grey, the disordered region in red and the VWA domain in pink. The 'finger' structure of the VWA domain located in the central pore of the NorQ hexamer is marked.*

***Table S2: AlphaFold multimer (AF-m) prediction datasets of this study and their corresponding pTM scores.***

| **Model** | **Average pTM** | **used for figure** | **Description** | **# models obtained** | **AF-m release** |
|---|---|---|---|---|---|
| NorD_Nterm-NorQ | 0.39 | not shown | N-terminal domain of NorD with the NorQ hexamer | 5 | v2.1.1 |
| NorD_VWA-NorQ | 0.49 | Figure 7d | C-terminal (VWA domain) of NorD and the NorQ hexamer | 5 | v2.1.1 |
| NorD-NorQx1-split | 0.69 | Figure 7c | N-terminal and C-terminal domains of NorD, without disordered domain, with NorQ monomer | 5 | v2.1.1 |
| NorD-NorQ_v2 | 0.41 | Figure S9 | NorD monomer plus NorQ hexamer | 5 | v2.2.0 |
| NorD-NorQx1 | 0.70 | Figure 7b | NorD and NorQ monomer each | 1 | v2.2.0 |

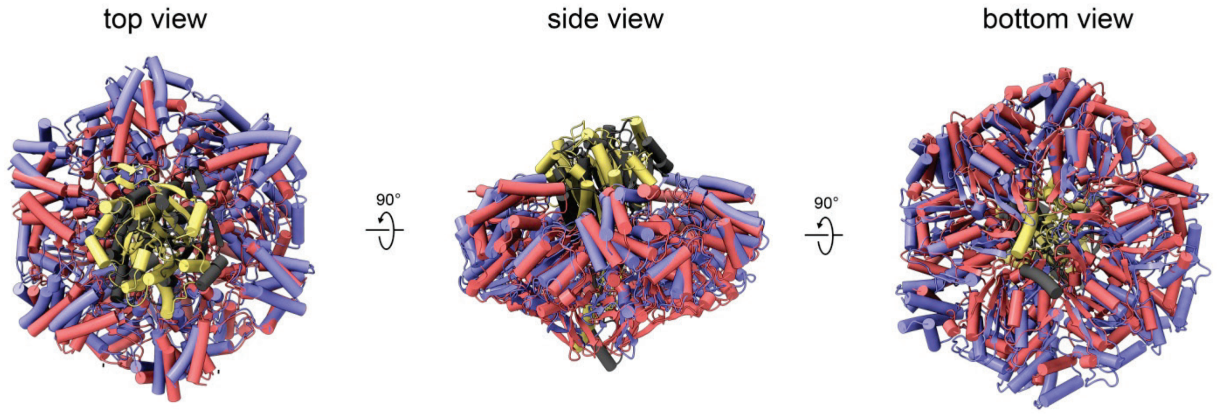


***Figure S10: Superposition of the AlphaFold prediction of NorQ$_6$VWA$_1$ (Fig 7d) and the fitted model based on our cryoEM map of the NorQD complex (Fig. 8a).*** *The NorQ$_6$VWA$_1$ AlphaFold model is shown in red (NorQ) and yellow (VWA) and our fitted model is in blue (NorQ) and black (VWA).*

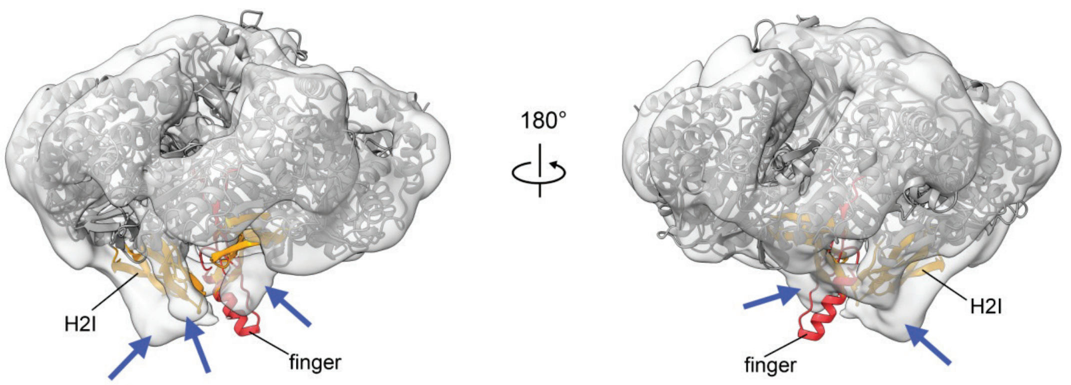


***Figure S11**: **Extra densities in the NorQ$^{WB}$D cryo-EM map. Shown is the modeled complex of the NorQ hexamer together with NorD VWA domain as fitted into the cryo-EM map (see main text Fig. 8).** Extra densities that were not well fitted on the convex side of the NorQ ring are marked with blue arrows. The extra densities are in close proximity to the H2Is of the NorQ protomers (yellow) and the 'finger' structure (red) protruding from the NorD VWA domain.*

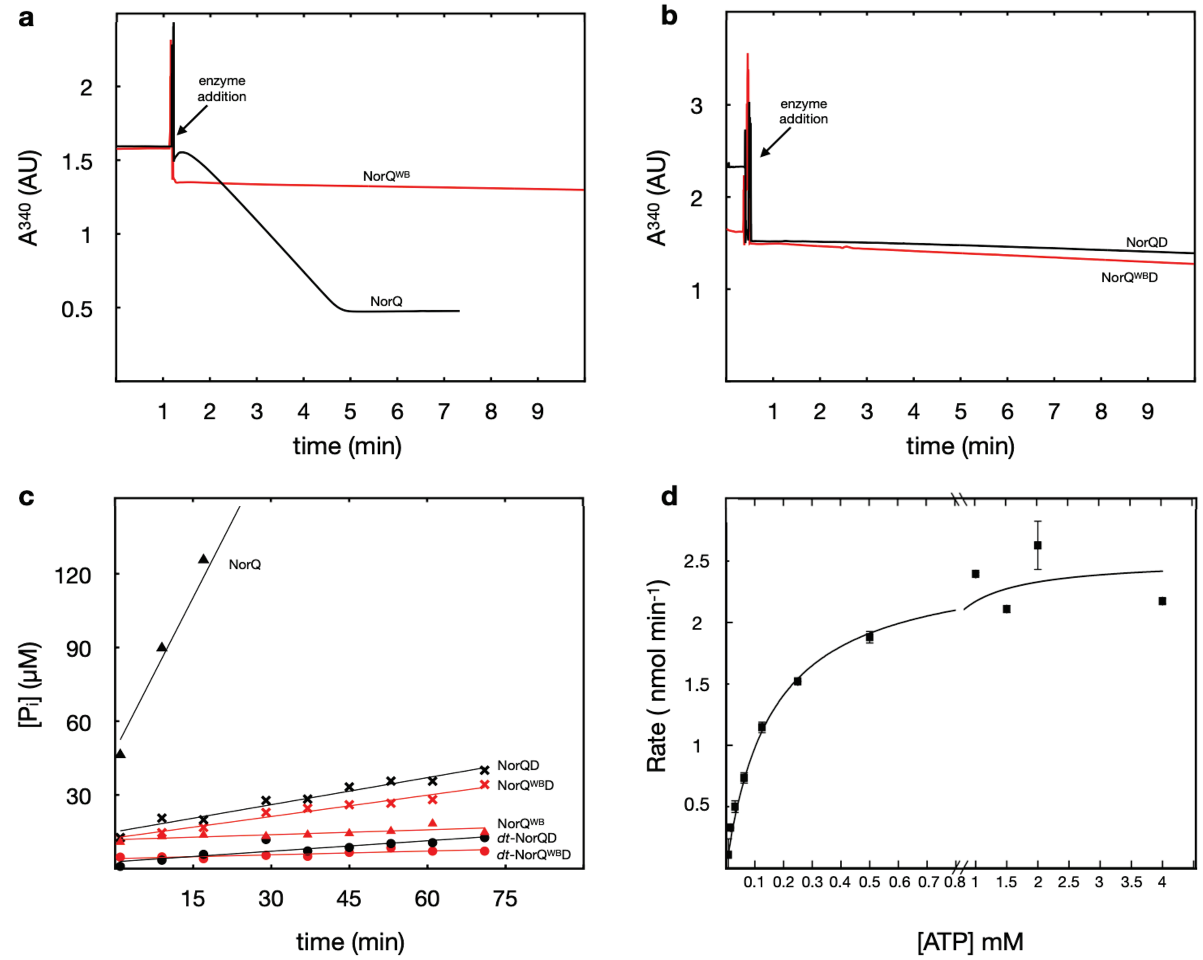


***Figure S12: ATPase activity measurements.*** *a) NorQ (black) and NorQ$^{WB}$ (red) in the NADH-coupled assay. b) NorQD (black) and NorQ$^{WB}$D (red) in the NADH-coupled assay. The reaction mixture contained 50 mM TRIS/HCl, pH 8.0, 150 mM NaCl, 15 mM $MgCl_2$, 2.5 mM ATP, 1 mM phosphoenolpyruvate, 0.3 mM NADH, 12 U/mL pyruvate kinase, 12 U/mL lactate dehydrogenase. c) Linear regressions for data from the malachite green end-point assay for NorQ (black triangles), NorQ$^{WB}$ (red triangles), NorQD (black crosses), NorQ$^{WB}$D (red crosses), double tagged NorQD (black circles) and double tagged NorQ$^{WB}$D (red circles). The initial reaction mixture contained 50 mM TRIS/HCl, pH 8, 150 mM NaCl, 10% (v/v) glycerol, 20 mM $MgCl_2$, 2 mM ATP. Data shown in A-C are example traces. d) Michaelis-Menten kinetics for NorQ (0.15 mg/ml). The data (shown as the average ± SD, n=2) was fitted to a $K_m$=0.16±0.02 mM, $V_{max}$=2.5±0.1 nmol min$^{-1}$ and a Hill coefficient = 1. The corresponding $k_{cat}$ is 20±1 ATP min$^{-1}$ hexameric complex$^{-1}$.*

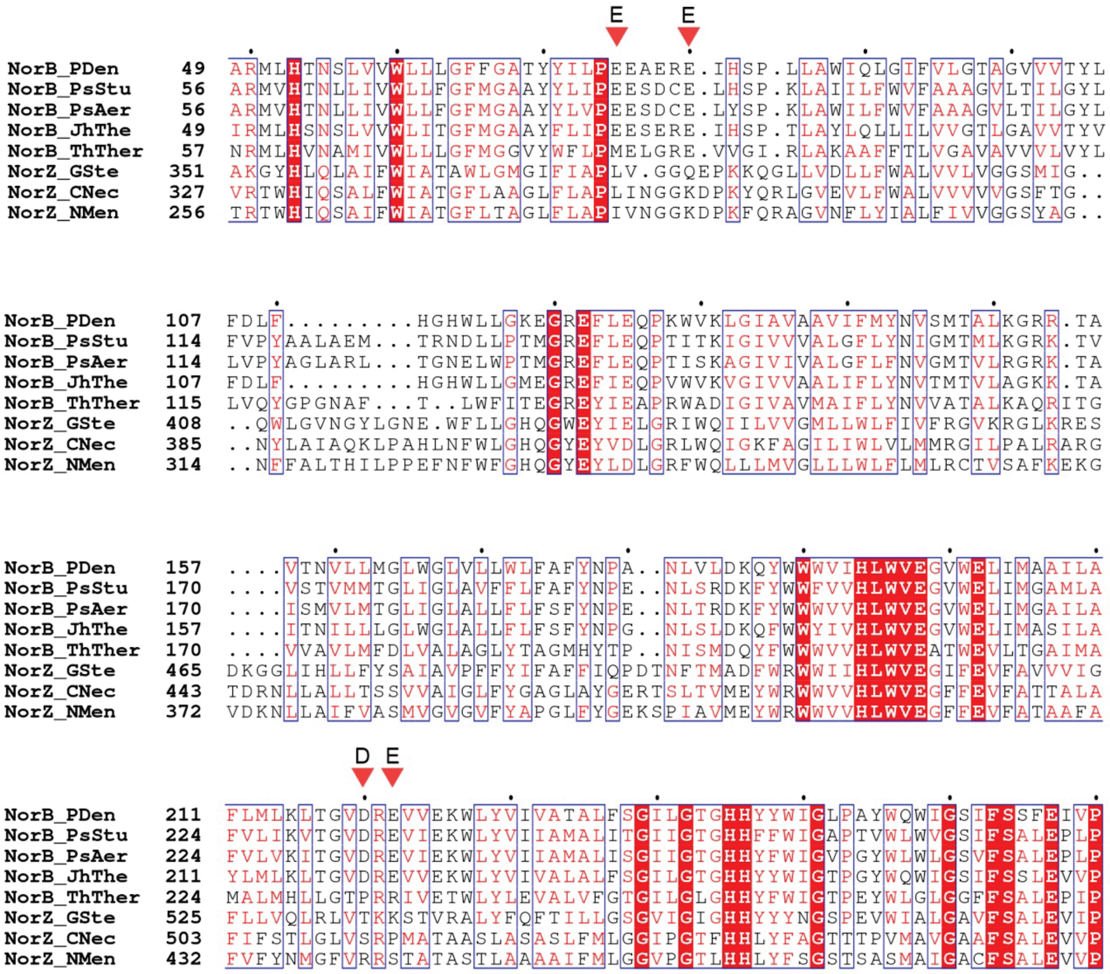


***Figure S13: Sequence alignment of the NorB subunit of cNOR with NorZ (qNOR) from various organisms.*** *We included cNOR from P. denitrificans (PDen; Uniprot: Q51663), Ps. stutzeri (PsStu; Uniprot: P98008), Ps. aeruginosa (PsAer; Uniprot: A0A072ZET2), Jhaorihella thermophila (JhThe; NCBI: WP_104006941.1) and Thermus thermophilus (ThTher; Uniprot: D5GU63). Sequences of qNOR are from G. stearothermophilus (GSte; B3Y963), Cupriavidus necator (CNec; O30375) and Neisseria meningitides (NMen; Q9JPL2). The acidic residues that were found conserved on the cytoplasmic surface of cNOR (excluding the T. thermophilus cNOR) but not in qNOR and that were mutated in this study are marked by red triangles. The sequence alignment was made using Clustal Omega (3) and ESPpript 3.0 (4).*

***Table S3: Re-activation experiments with cNOR using crude cell extracts***. *The data are presented as the average value (± SD, n=2).*

| ***Genes present (soluble fraction)*** | ***Genes present (membrane fraction)*** | ***Activity of purified cNOR (%)*** | ***Non-heme Fe content (%)*** |
|---|---|---|---|
| *norCBQDEF* | *norCBQDEF* | *100 ± 40* | *100 ± 10* |
| *norCBQDEF* | *norCB(ΔQDEF)* | *19 ± 1* | *21 ± 1* |
| *norCB(ΔQDEF)* | *norCB(ΔQDEF)* | *7.8 ± 0.3* | *10 ± 1* |

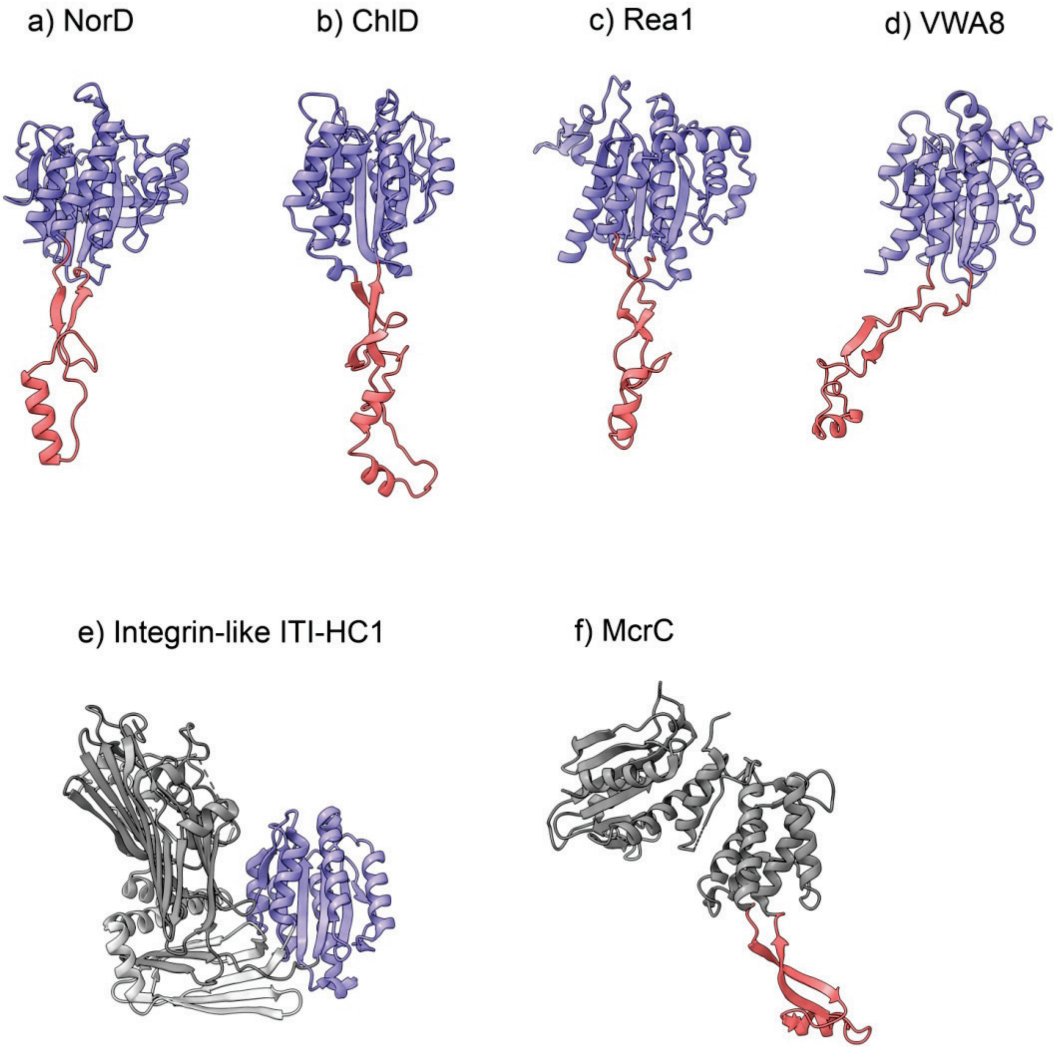


***Figure S14**: **Predicted VWA domain of NorD (a) in comparison to several eukaryotic VWA domains (b-e) to McrC (f).** The predicted VWA domains (a-d) are all models produced by AlphaFold.VWA domains in blue and finger-like protrusions in red. Remaining structural elements are colored in grey. a) NorD from P. denitrificans (Uniprot: Q51665), b) ChlD from Arabidopsis thaliana (Uniprot: Q9SJE1), c) Rea1 from Saccharomyces cerevisiae (Uniprot: Q12019), d) human VWA8 (Uniprot: A3KMH1), e) crystal structure of a human ITI-HC1 monomer (PDB: 6FPY) (1). f) cryo-EM structure of McrC from E. coli (Uniprot: P15006, PDB: 6HZ4 (8)).*

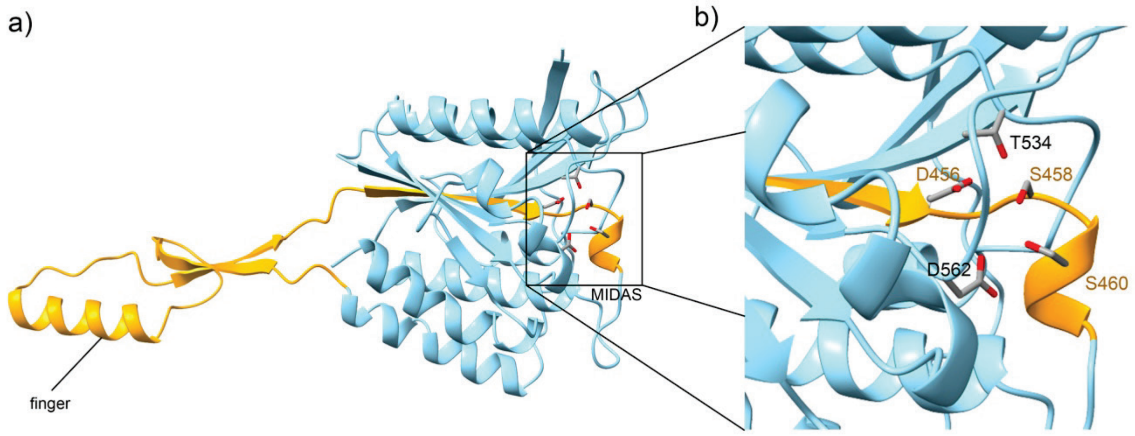


***Figure S15: Structural connection between the 'finger' and the MIDAS in the VWA domain of the AlphaFold prediction model of NorD.*** *a) The 'finger' and MIDAS are positioned at opposite sides of the VWA domain, but a single β-strand (orange) connects the 'finger' to the MIDAS residues D456, S458 and S460 located in the VWA core (cyan). b) Close-up of the MIDAS, with the residues indirectly connected to the 'finger' via a single β-strand labeled in orange.*

### Table S2: Primers used in this study.

| function | forward primer | reverse primer |
|---|---|---|
| N-terminal His tag NorQ in pNOREX | CGGAGTAAGCCATGCATCATCATCATCATCATAACGCGCATGTGAAAACCCAAGGG | CACATGCGCGTTATGATGATGATGATCATGCATGGCTTACTCCGCAGCCTCC |
| N-terminal His tag NorD in pNOREX | GGGAAGCGCCATGCATCATCATCATCATCATGGTCTGGACCTGGAACCC | GGTCCAGACCATGATGATGATGATGATGCATGGCGCTTCCCCTCAGCC |
| N-terminal His tag NorQ in pET21 | GAGATGCGCGTTGCTAGCATGATGATGATGATGATGCATATGTATATCTCCTTCTTAAAGTTAAAC | CACATGCGCGTTGCTAGCATGATGATCATGATGATGCATATGTATATCTCCTTCTTAAAGTTAAAC |
| N-terminal His tag NorD in pET21 | CGCCATGCATCATCATCATCATCATTCTAGAGGTCTGGACCTGGAACCCTGGGAGC | CCAGACCTCTAGAATGATGATGATGATGATGCATGGCGCTTCCCCTCAGCCGAA |
| Cloning NorQ into pETDuet-1 | TGAGATCTCAACGCGCATGTGAAAACCC | TGATCTCGAGGCCGAAGATCGCGGCG |
| Cloning NorD into pETDuet-1 | ATCGAATTCGGGTCTGGACCTGGAACCC | GATAAGCTTTCACGCCACCAGTTGCCGATAG |
| Exchange of S-tag for Strep tag II in pETDuet-1 | ACTGCTCGAGAGCGCTTGGAGCCACCCGCAGTTCGAAAAATGACCATGGTTAATTAATGA | TCATTAATTAACCATGGTCATTTTTCGAACTGCGGGTGGCTCCAAGCGCTCTCCAGCAGT |
| Amplifying NorQ and adding restriction sites | AGAGCTAGCAACGCGCATGTGAAAACCC | TGTCTGAATTCCTTCCCCTCAGCCGAAGATC |
| Amplifying NorD and adding RBS and restriction sites | AGCGAATTCAAGAAGGAGATATACATATGCATCATCATCATCATCATTCTAGAGG | TCTAAGCTTTCACGCCACCAGTTGCC |
| Walker B mutation in NorQ (E109Q) | CTCGACCAGGTGGTCGAGGCGCGCAAGGACG | GACCACCTGGTCGAGATAGCAGATCGCGCCCTCG |
| Walker A mutation in NorQ (K48A) | GCGGCGCGACCCGCTTCGTCGCCCATATGG | GCGGGTCGCGCCGCAGCCGGTCGGGCCTTTCAGC |
| T534V mutation in NorD | GCGCTTCTATGTGCGGCTTGGCGCGG | CCAAGCCGCACATAGAAGCGCGGCCTCAGC |
| D562N mutation in NorD | GGTCATCACCAACGGCAAGCCCAACGACCTGG | GCTTGCCGTTGGTGATGACCAGAAGCAGCCG |
| E75A mutation in NorB | GCCCGCAGAGGCCGAGCGCGAGATCCATTCGCC | GCCTCTGCGGGCAGGATGTAATAGGTCGCGCCG |
| E78A mutation in NorB | GGCCGCGCGAGTCCATTCGCCGCTGC | GCGCGCGGCCTCTTCGGGCAGGATGTAATAGGTC |
| Double mutation E75A and E78A in NorB | CCCGCAGAGGCAGCGCGCGAGATCCATTCGCCGCTG | GCGCGCTGCCTCTGCGGGCAGGATGTAATAGGTCGC |
| D220A mutation in NorB | CGTGGCCCGCGAGGTGGTCGAGAAATGGCTTTACGT | CGCGGGCCACGCCGGTCAGCTTGAGCATCA |
